# stLearn: integrating spatial location, tissue morphology and gene expression to find cell types, cell-cell interactions and spatial trajectories within undissociated tissues

**DOI:** 10.1101/2020.05.31.125658

**Authors:** Duy Pham, Xiao Tan, Jun Xu, Laura F. Grice, Pui Yeng Lam, Arti Raghubar, Jana Vukovic, Marc J. Ruitenberg, Quan Nguyen

## Abstract

Spatial Transcriptomics is an emerging technology that adds spatial dimensionality and tissue morphology to the genome-wide transcriptional profile of cells in an undissociated tissue. Integrating these three types of data creates a vast potential for deciphering novel biology of cell types in their native morphological context. Here we developed innovative integrative analysis approaches to utilise all three data types to first find cell types, then reconstruct cell type evolution within a tissue, and search for tissue regions with high cell-to-cell interactions. First, for normalisation of gene expression, we compute a distance measure using morphological similarity and neighbourhood smoothing. The normalised data is then used to find clusters that represent transcriptional profiles of specific cell types and cellular phenotypes. Clusters are further sub-clustered if cells are spatially separated. Analysing anatomical regions in three mouse brain sections and 12 human brain datasets, we found the spatial clustering method more accurate and sensitive than other methods. Second, we introduce a method to calculate transcriptional states by pseudo-space-time (PST) distance. PST distance is a function of physical distance (spatial distance) and gene expression distance (pseudotime distance) to estimate the pairwise similarity between transcriptional profiles among cells within a tissue. We reconstruct spatial transition gradients within and between cell types that are connected locally within a cluster, or globally between clusters, by a directed minimum spanning tree optimisation approach for PST distance. The PST algorithm could model spatial transition from non-invasive to invasive cells within a breast cancer dataset. Third, we utilise spatial information and gene expression profiles to identify locations in the tissue where there is both high ligand-receptor interaction activity and diverse cell type co-localisation. These tissue locations are predicted to be hotspots where cell-cell interactions are more likely to occur. We detected tissue regions and ligand-receptor pairs significantly enriched compared to background distribution across a breast cancer tissue. Together, these three algorithms, implemented in a comprehensive Python software stLearn, allow for the elucidation of biological processes within healthy and diseased tissues.

## Background

Recent times have seen breakthroughs in the discovery of new cell types and deepened understanding of how they affect health or respond to changes in their microenvironments, through the advent of single-cell RNA sequencing (scRNAseq). scRNAseq is an ultra-sensitive and high-throughput technology to identify cell types and transcriptional states to individual cell resolution^1^. However, tissues represent enormously complex cellular ecosystems characterised by local cell type composition, cell distribution, and cell-cell interactions^2–4^. The nature of these features at any given place and time within a tissue are critical determinants of tissue development, homeostasis, repair, and responses to environmental signalling^2, 5^. Current knowledge about cell types still lacks crucial information about how these cell types coexist within their native tissue ecosystems in healthy and diseased states^6–8^.

Spatial transcriptomics (ST) is rapidly emerging as the “next generation” of scRNAseq^9^. ST can unbiasedly profile transcriptome-wide gene expression and does not require tissue dissociation, hence retaining spatial information. Experimental methods to generate ST data are being established, using platforms such as Visium from 10X Genomics^10^, NanoString DSP from NanoString^11^, seqFISH^12^, MERFISH^13^, and Slide-seq2^14^. Laboratories around the world have begun releasing data^15^, with the number of public ST datasets highly expected to surpass the current number of 300 sub-datasets^15^ in the near future. In this work, we focus on one of the first and most established ST technologies^16^, which is now widely available as a Visium platform by 10x Genomics. For the same tissue section, the technology provides users with gene expression data, spatial distance information, and a hematoxylin and eosin (H&E)-stained microscopy image with tissue morphology information. stLearn can also be used to analyse other ST data types, such as slide-seq^14^ and MERFISH^13^.

Finding patterns in spatial gene expression data is one of the great challenges in single-cell-omics data science today^8^. As with scRNAseq, ST analysis tool development has lagged behind rapid technological and data generation advances. To find cell types, existing clustering algorithms for ST data use only gene expression information, such as in Spaniel^17^, Seurat (V3.2)^18^, spatialLIBD^10^, and SpatialCpie^19^. Current available methods to find differentially expressed genes^20, 21^ or spatially variable genes^22^, can integrate gene expression and spatial distance information but not tissue-image information. However, morphology and gene expression are strongly linked^23^. Our previous work pioneering the combinatorial use of imaging pixel information with gene expression data in a neural network model demonstrated that morphology and gene expression are strongly linked and that using either imaging or gene expression data alone were less accurate (lower AUC and prediction accuracy) at predicting cell types and disease stages compared to models combining both data information^24^.

In this study, we present a significant advancements in the integrating of gene expression measurements with spatial distance information and tissue morphological information which will enable researchers to make optimal use of ST data. Our methods described here addresses three major research areas: the identification cell types, the reconstruction of cell trajectories, the study of cell-cell interactions within a morphologically-intact tissue section. We hypothesised that the inclusion of spatial information and cellular morphological data can address the challenges in these three research areas with higher sensitivity and accuracy.

In vitro cell state trajectories can be built from snapshots of single-cell transcriptional profiles^25^. Traditionally, most statistical models to study cell progressions – for instance, across cell differentiation, cancer evolution, or injury progression – have not considered spatial structure^8, 26^. To our knowledge there is not yet a method that reconstructs biological trajectories within a tissue. In this work, we adapt the concept of pseudotime^27^ and introduce a pseudo-space-time (PST) algorithm, where gene expression and spatial distance are integrated to model transcriptional relationships between cell types within a tissue.

Another advantage of spatial data is the ability to study cell-cell interactions (CCI) to discover where and how neighbouring cells affect transcriptional networks in their native tissue. Existing methods to model cell-cell interactions are based on testing for enrichment in ligand-receptor (L-R) expression from bulk RNAseq or scRNAseq data^28, 29^. Utilising spatial distance and cell type distribution within the tissue, we present a novel method for detecting CCI hotspots automatically and unbiasedly across the whole tissue.

Together, this work introduces three main algorithms that allow the integration of spatial location, tissue morphology and gene expression, to answer significant biological questions about cell type identification, transcriptional connectedness between cell types, and cell-cell interactions. These algorithms are implemented in a user-friendly Python-based software, stLearn. In one streamlined package, researchers can mine ST data from raw inputs to in-depth downstream analysis and generate intuitive graphics to visualise results.

## Results

### 1. Overview of the stLearn analysis workflow

The results and methodological background for main analyses and algorithms are described in subsequent sections. Briefly, we present a comprehensive analysis workflow comprising novel approaches to key tasks in ST data analysis (Fig. 1), including: *1)* normalisation (by neighbourhood smoothing and morphological adjustment); *2)* clustering (using normalised data and graph-based clustering to find clusters that are spatially and transcriptionally defined; *3)* spatial trajectory analysis (PST for finding local and global relationships between cell types); *4)* cell-cell interactions and microenvironment detection, and *5)* other common tasks such as dimensionality reduction including principal component analysis (PCA), Uniform Manifold Approximation and Projection (UMAP), Independent Component Analysis (ICA), Factor Analysis (FA), and Diffusion Pseudotime (DP), and visualisation such as plotting to add to the tissue image the following information: gene expression, cluster labels, sub-cluster lebels, microenvironments, and in vivo trajectories.

**Figure 1.**
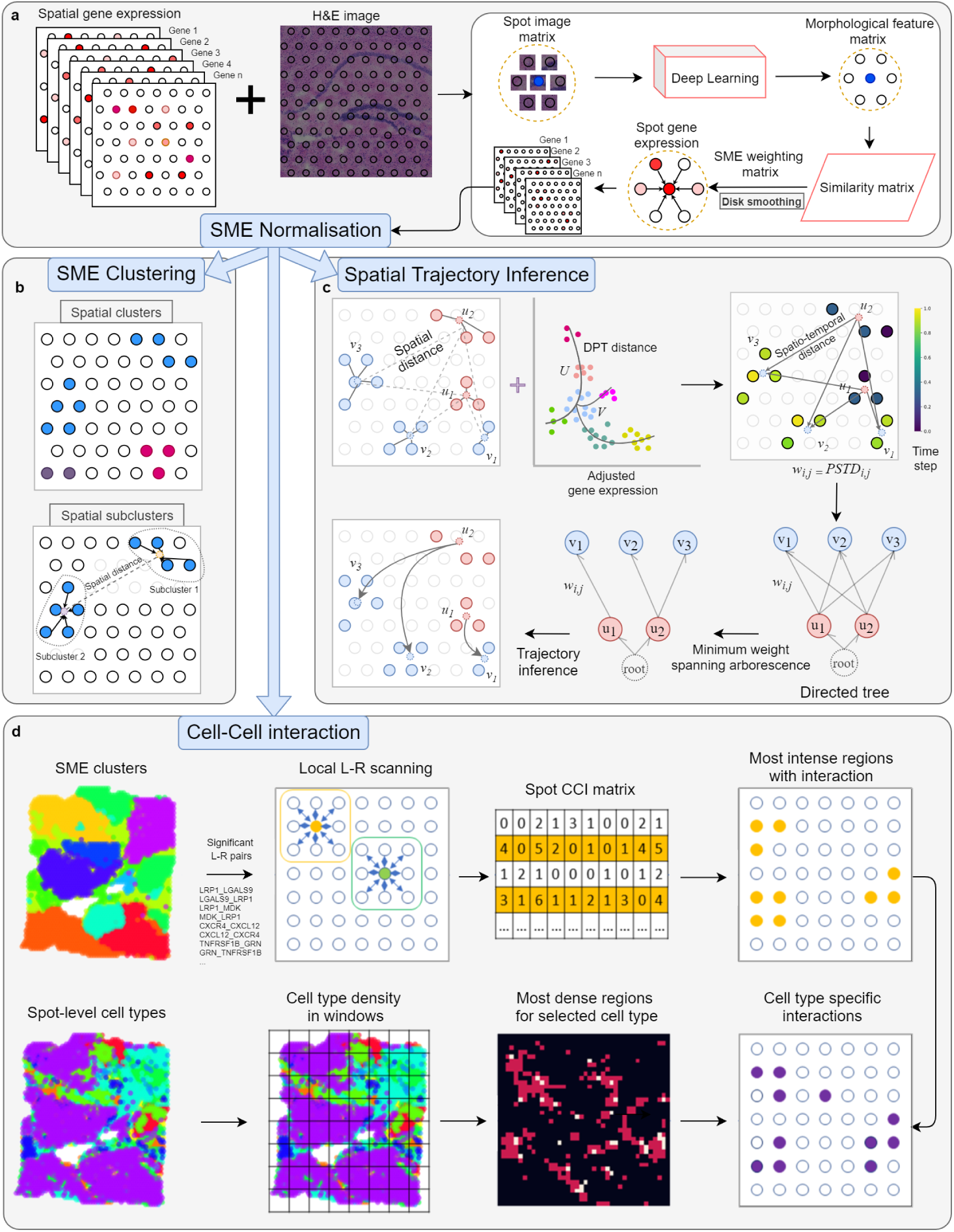
Innovative spatial analysis approaches implemented in stLearn. **a**, Schematic diagram of the processes involved in Spatial Morphological gene Expression Normalisation (SME Normalisation). Briefly, a within-tissue normalisation step that adjusts gene expression values by spatial location and morphological similarity. ST arrays are represented as large squares with many coloured spots, with red shading to indicate hypothetical gene expression intensity. In each ST array, thousands of genes are measured for each spot. Neighbouring spots within a radius *d* in a tissue are used for normalisation. Morphological data from the H&E image in the form of a three-dimensional array (height × width × colour) is processed by a deep learning model to form a two-dimensional feature matrix. The matrix is used to compute morphological similarity (distance) between neighbouring spots. Each gene within a spot is re-assigned a normalised expression value representing the mean expression of all neighbouring spots weighted by the morphological similarity (SME expression values). After SME normalization, the following analysis were carried out: SME clustering, spatial trajectory inference and cell-cell-interaction analysis. **b**, Clustering for cell type identification. SME clustering is performed based on SME normalised expression values, through dimensionality reduction and graph-based community detection. The SME-based cluster labels are plotted to the H&E tissue image. Here, cluster labels are represented by coloured spots (blue, purple and red dots) mapped to their spatial locations in the ST arrays. If spots of a cluster are spatially separated into two or more locations in the ST array (such as the blue cluster shown here), the initial cluster can be split into sub-clusters. **c**, Spatial trajectory analysis. Physical (spatial) distance is calculated between centroid coordinates of SME clusters *U* and *V* with sub-clusters (*u*_1_, *u*_2_) and (*v*_1_, *v*_2_, *v*_3_). Diffusion pseudotime (DPT) is calculated based on gene expression. These measures are combined to produce pseudo-space-time distance (PSTD), which is used to form a directed graph that can be optimised by minimum spanning rooted tree for trajectory inference. **d**, Cell-cell interaction analysis. Ligand-Receptor co-expression enrichment among neighbour spots and cell-type density enrichment are combined to find hotspots within a tissue, where interactions between cell types are most likely to occur.

### 2. SME normalisation

SME Normalisation (Spatial Morphological gene Expression normalisation) is a within-tissue normalisation strategy that uses neighbourhood information (spatial location) and morphological distance to normalise gene expression data. SME normalisation represents the first step in the stLearn workflow, where disk smoothing is used to adjust the expression value in each spot based on the surrounding proximal spots within a given radius *d* (Fig. 1a). Disk smoothing makes the key assumption that two spots that are nearby together or that morphologically resemble each other will have more similar gene expression compared to more distant or morphologically distinct spots (Fig. 1a). This assumption is supported by previous studies demonstrating correlation between gene expression and morphology^23^, or have reported genes with localised expression patterns^21^. The H&E image of the tissue section is split into small tiles, each containing one spot. Feature vectors are extracted from these pixel image tiles by a transfer learning deep neural network model (Fig. 1a)^30^. Then, for a given centre spot and its neighbouring spots, a morphological similarity measure is calculated by pairwise cosine distance of feature vectors between centre spot *S* and its neighbour spots *S*_*i*_. In this way, the expression value for each gene in the centre spot is calculated as the mean (by default, or median) of morphological similarity-weighted expression values of *n* neighbouring spots. By using information from neighbouring spots, we reduce the technical limitation of poor detection of lowly expressed genes, which appear as zeros in the expression matrix, an issue often referred to as dropouts. A zero-expression value for a gene in one spot occurs only if all of its neighbouring spots also have an expression value that is zero. SME normalisation is highly valued to analyse noisy spatial sequencing data, because information from the high resolution H&E image allows the inherent sequencing limitations of technical noise and detection sensitivity to be adjusted. To our knowledge, this novel algorithm is the first to use imaging information (cellular morpholoy) to adjust gene expression.

### 3. stLearn spatial clustering (SMEclust)

Using SME normalised data, stLearn performs unsupervised clustering to group similar spots into clusters, and to find sub-clustering options based on spatial segregation of clusters in the tissue. This stLearn function is called SMEclust. Using normalised expression values, stLearn performs a two-step spatial clustering procedure to segregate cell types in a tissue (Fig. 1c, 2). For the first step, stLearn implements a standard Louvain clustering procedure as applied to scRNAseq data in scanpy^31^ or Seurat^18^. The SME normalised matrix is the input for linear PCA dimensionality reduction, followed by non-linear UMAP embedding, and k-nearest neighbour (kNN) graph construction. Louvain clustering^32^ or k-means clustering is then applied to the graph adjacency matrix from kNN. In the second step, spatial information is used to find sub-clusters from broad clusters which are spread across two or more spatially separated locations. Spatial coordinate information for of each spot is used for performing a two-dimensional k-d tree neighbour search (Fig. 1c). We found that SMEclust had higher clustering performance compared to other methods, as described in two case studies below (Fig. 2, Supplementary Figs. 1-4).

**Figure 2.**
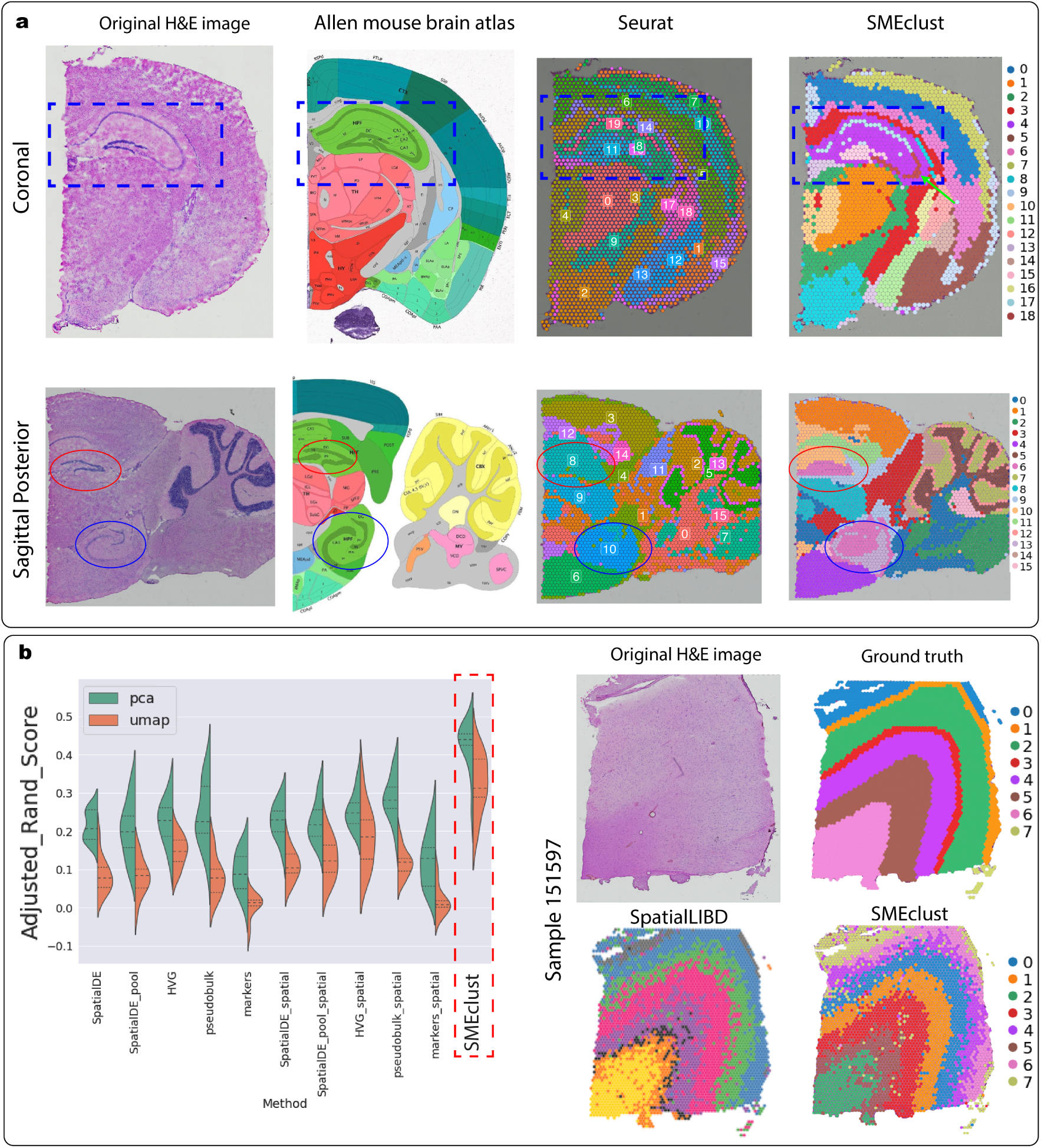
Spatial clustering (SMEclust) in mouse and human brain tissue sections. **a**, Clustering of mouse brain tissue coronal (top) and sagittal posterior (bottom) sections. The H&E image generated from the data (first column) and the corresponding anatomical Allen Mouse Brain Atlas (second column) are shown. From identical input, clustering results by the Seurat spatial pipeline (third column) and the SMEclust method (fourth column) were compared. SMEclust detected anatomical regions that Seurat missed, for example the CA3 region in the coronal section (Cluster 18) (blue box), and the dentate gyrus in the sagittal section (blue and red cicles). Similar comparisons for the sagittal anterior section are shown in Supplementary Fig. 1. **b**, Clustering analysis for human dorsolateral prefrontal cortex data^10^. For each of 12 tissue sections, we compared SMEclust to spatialLIBD run with 20 parameter combinations (10 ways of selecting gene features for clustering and two ways for dimensionality reduction). SMEclust and spatialLIBD results were compared to the ground truth data reported in the original paper to compute the adjusted Rand index for each sample, a parameter for pairwise comparison of two clustering results. Example clustering results for tissue section 151673 are shown here (right). Clustering results for two additional tissue sections are shown in Supplementary Fig. 3, and additional statistical test for comparing adjusted rand index values are shown in Supplementary Fig. 4.

#### Case study 1

We tested SMEclust on mouse brain datasets, and compared our results to the brain anatomical reference annotations from the Allen Mouse Brain Atlas^33^ were considered as ground truth (Fig. 2, Supplementary Fig. 1, 2). Firstly, we compared the abilities of SMEclust and Seurat (V3.2) to detect anatomical cell types in mouse brain coronal, sagittal anterior and sagittal posterior tissue sections (Fig. 2a, Supplementary Fig. 1). Compared to the Allen Mouse Brain Atlas, SMEclust could detect the CA3 (cornu ammonis 3, shown as cluster 18) area of the hippocampus region, while Seurat could not (in both cases for coronal and sagittal posterior sections; Fig. 1a, Supplementary Fig. 1a). SMEclust could also split cerebral cortex (CTX) into six layers, an improvement over Seurat which could only detect four layers (sagittal anterior section; Supplementary Fig. 1b). Secondly, we applied a common coordinate framework^33, 34^ to accurately overlay the three-dimensional reference brain from the Allen Mouse Brain Atlas onto the corresponding brain tissue section from our data (Supplementary Fig. 2). Consistent trends in SMEclust and Seurat capabilities were observed for all three mouse brain tissue sections (Supplementary Fig. 1, 2).

#### Case study 2

We also tested SMEclust on ST data from all twelve human brain tissue sections^10^ and compared the results to those generated with another ST clustering algorithm, SpatialLIBD, which was applied in the original publication of this data. We tested five options that SpatialLIBD used to select gene markers: SpatialDE genes, SpatialDE pooled genes, highly variable genes (HVG), pseudo-bulk, and known markers. For each of these five options, we tested the effects of inclusion and exclusion of spatial information. Clustering was then performed in both PCA and UMAP spaces. Overall, this resulted in 20 different combinations of SpatialLIBD parameters. We found that SMEclust outperformed all twenty SpatialLIBD parameter combinations for generating data more consistent with the ground truth (Fig. 2b). We also observed this trend in all other human brain tissue sections tested (Supplementary Fig. 3, 4).

### 4. stLearn spatial trajectory inference

To discover *in vivo* processes happening within an intact tissue – for example, to answer which cancer cell or clone appeared first, or how a cancer evolved – we developed an algorithm called pseudo-space-time (PST) trajectory analysis. PST is an extension of the pseudotime concept commonly applied in scRNAseq data analysis^27^, and is designed to detect biological processes that can be inferred from gradient changes in transcriptional states across a tissue. With PST, evolutionary trajectories are reconstructed based on transcriptome profiles and the spatial context of cells within a tissue. Following SMEclust analysis, we implement the PST algorithm to find relationships between and within clusters at local (i.e. relationships between sub-clusters of a single cluster) and global (i.e. relationships between clusters) levels in a tissue. First, trajectory analysis using PAGA^35^ based on whole-tissue SME-normalised gene expression data is used to find connections within clusters. Next, pseudotime is computed via the diffusion pseudotime (DPT) method^36^ which is a robust trajectory inference method. The DPT method measures cell-to-cell transitions using diffusion-like random walks. For the direction of the trajectory, a root can be defined by the user depending on which biological processes are being investigated in the tissue. We then compute pseudo-space-time distance (PSTD), which combines both gene expression values and physical distance (see Methods section). Given two sets of spots *U* and *V* (belonging to two cell types or two clusters) with their corresponding centroids *u* and *v*, we apply a formula to combine physical distance with gene expression distance, with *ω* as a user-specified weighting factor reflecting the balance between these two values. With the adjusted PSTD matrix, we construct a directed graph, and then optimise the graph using the directed minimum spanning tree algorithm to find the shortest, rooted tree and branches (trajectories) that connect nodes in the graph^37^. The PST algorithm is described in detail in the Method section.

#### Case study 3

As a proof of concept, we aimed to explore tumour evolution from ductal carcinoma *in situ* (DCIS) to invasive ductal carcinoma (IDC), known as DCIS-IDC progression. Human Breast Cancer (Block A Section 1) spatial data, publicly available from 10x Genomics, was used. This dataset consists of 3,813 spots within the tissue area and 22,968 genes with a median of 5,394 genes per spot. SMEclust predicted 11 clusters (Supplementary Fig. 6b) and their corresponding sub-clusters (Supplementary Fig. 6d). Based on H&E morphology and differential expression of marker genes, we identified one DCIS cluster comprised of three sub-clusters (sub-clusters 6, 11 and 14), and one IDC cluster comprised of seven sub-clusters (sub-clusters 9, 15, 19-21, 24 and 39)(pink and sky blue, respectively, in Supplementary Fig. 6b).

To model cancer progression from DCIS to IDC, the PST algorithm was applied to find the spatial and transcriptional connections between the DCIS and IDC sub-clusters. We also built a local spatial trajectory inference (Supplementary Fig. 6e) (DCIS subclonal evolution) by constructing a PSTD matrix (Supplementary Fig. 6a, 6c) as described in the Methods section. Trajectory optimisation is processed by using a minimum directed spanning tree approach. As a result, we differentiated three lineages of tumour evolution (Fig. 3a, 3c), whereas only a single unbranching lineage is detected if gene expression alone is used (pseudotime analysis) (Fig. 3b). In addition to spatial visualisation, stLearn allows visualisation of a hierarchical tree of cancer cell type evolution (Fig. 3c) to facilitate interpretation of results. Compared to the single DCIS-to-IDC progression pathway detected with pseudotime analysis, our PST algorithm allowed us to detect three DCIS-to-IDC clades (i.e. clade 1: 6-19, 6-20; clade 2: 11-15, 11-21, 11-24; clade 3: 14-39; Fig. 3c), reflecting the branched evolution inherent to cancer progression^38, 39^.

**Figure 3.**
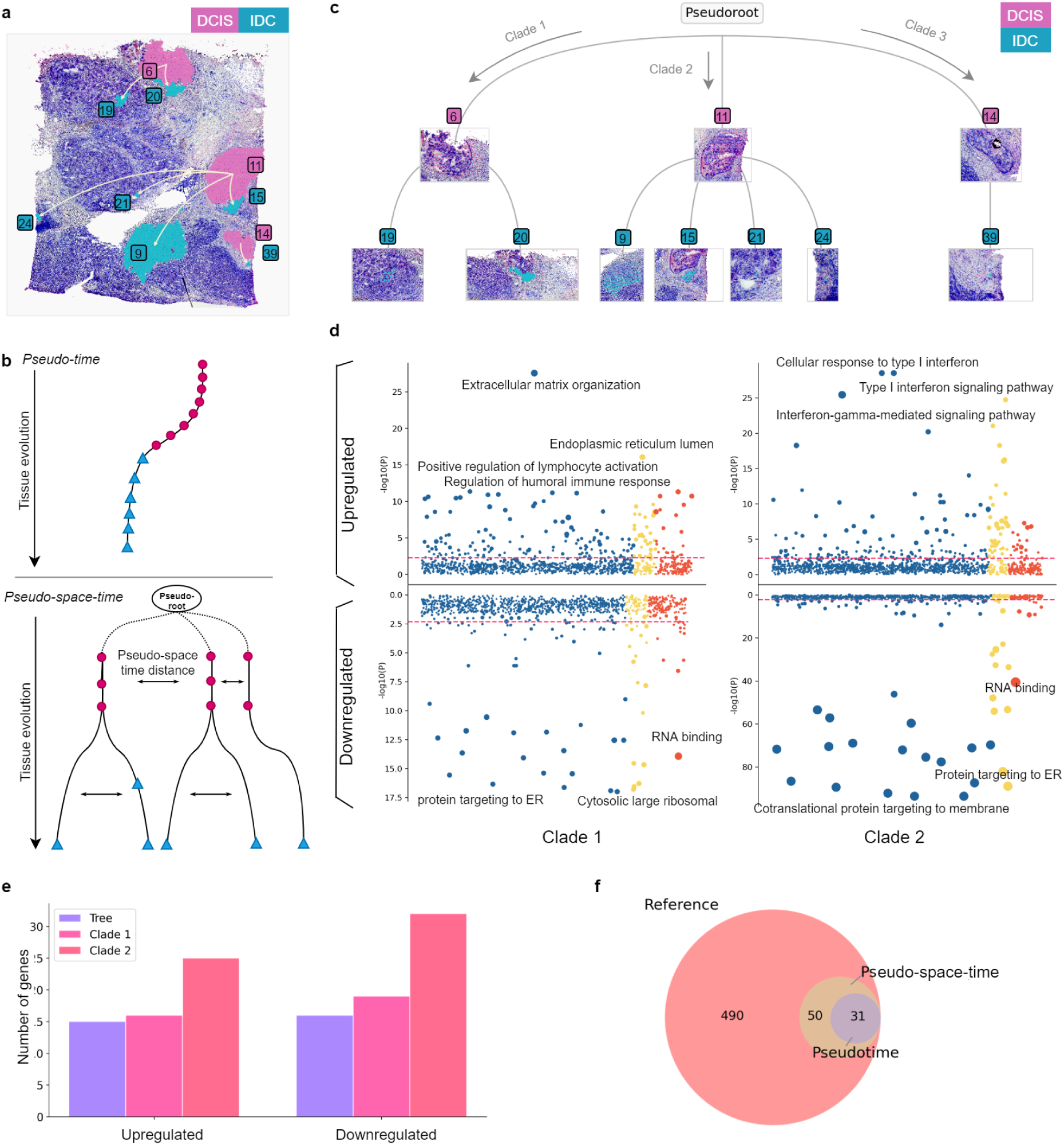
Pseudo-space-time (PST) analysis of breast cancer tissue. **a**, Visualisation of DCIS-IDC (ductal carcinoma in situ – invasive ductal carcinoma) trajectories with tissue localisation. Pseudo-space-time distance (PSTD) calculations (*ω* = 0.5) led to the identification of three DCIS sub-clusters leading to multiple IDC sub-clusters(Clade 1: (6-19, 6-20), Clade 2: (11-9, 11-15, 11-21, 11-24) and Clade 3: (14-39). **b**, Hypothetical representation of the pseudotime and PST methods. Pseudotime analysis generates a single linear trajectory of tissue evolution. The PST algorithm uses the combination of spatial information and gene expression profiles in trajectory inference to identify location-specific cancer evolution in tumour tissue. The pseudo-root represents the unidentifiable mutated cell that is the common ancestor of all descendant lineages; PSTD (shown here as arrows) is the distance between two clusters or sub-clusters based on both spatial distance and DPT; red circles represent hypothetical cells from an earlier developmental stage (for example, DCIS), blue triangles represent some later developmental stage (such as IDC). **c**, PST tree plot with hierarchical layout showing the flow of cancer progress in three clades branching from three DCIS regions (pink) to multiple IDC sub-clusters (blue) **d**, Bubble plot showing −log10(P values) for Gene Ontology (GO) enrichment analysis on the top 100 upregulated and top 100 downregulated genes in clades 1 and 2. The dashed line represents a P-value cut-off of 0.05. GO categories are coloured: GO biological process (blue), GO cellular component (yellow) and GO molecular function (red). Bubble sizes reflect the number of detected genes which contributed to a corresponding pathway or component. **e**,**f**, A group bar chart and Venn diagram showing the number of cancer driver genes found by either the unbranching pseudotime estimate (“tree”) or PST method (“clade 1” and “clade 2”) based on Pearson correlation with a cut-off of 0.2. The cancer driver genes were retrieved from The Cancer Gene Census (CGC) catalogue.

We performed trajectory-based differential expression analysis which identified transition genes – that is, genes that increase or decrease their expression along the trajectory in a branch- or location-specific manner (Supplementary Fig. 9a). Thus, a gene is considered to be upregulated if its expression increases along the DCIS-to-IDC trajectory, and downregulated if its expression decreases along this trajectory. Then, gene set enrichment analysis based on gene ontology was used to find differences between the three clades (Fig. 3h). For clade 1, we detected changes in genes associated with extracellular matrix organisation *(COL18A1, SPARC, COL12A1, LOXL3)* – a key driving process for both cancer development and progression^40^. We also found immune response genes *IGHG3, IGHG4, IGHG1, IGHG2, IGKC, C3, CD81, C1QB* to be positively correlated with DPT (i.e. upregulated genes), suggesting possible immune responses to or involvement of immune signalling in cancer cell expansion. Genes that were downregulated (i.e. negatively correlated with DPT, with high expression in DCIS and low expression in IDC) included those involved in endoplasmic reticulum stress^41^, dysregulation of ribosome biogenesis^42^ and nonsense-mediated mRNA decay^43^. This observation is consistent with the common mechanism that cancer cells grow strong and fast inside the DCIS and expands to nearby locations, forming IDC, where immune response processes are triggered^44^. Unlike observations from clade 1, the evolutionary progress in clade 2 was more likely to be driven by type I interferon^45^, with top upregulated genes including *IFITM3, STAT1, IFI6, ISG15, IFI35, HLA-C, HLA-A, HLA-F, HLA-E* identified. Moreover, IDC sub-cluster 9 comprised markedly more cells than other sub-clusters, suggesting that clade 2 may represent the stronger immune signaling pathways in areas to which the invasive cancer is spreading^46^. Clade 3 appears to represent an early stage of cancer invasion, as we found few significant changes in gene expression across the DPT (Supplementary Fig. 7b). We therefore did not include clade 3 in subsequent analyses. Overall, enrichment analysis of transition genes detected by the PST method revealed a location-specific biological insight of branched evolution in cancer progression.

To further compare pseudotime and PST methods, we next examined the number of cancer driver genes detected using the aforementioned trajectory-based differential expression analysis. We examined trajectory differentiation-associated genes for three sets of spots: tree, clade 1 and clade 2. Tree is defined by all spots from DCIS and IDC clusters, thus representing the result of the pseudotime method. Clade 1 and 2 are the trajectories inferred by PST. Genes were identified based on the 572 cancer driver genes in the Cancer Gene Census (CGC) catalogue. Based on Pearson correlation with DPT, targeted analysis of clades 1 and 2 identified more significant cancer driver genes (16 and 25 upregulated genes; 25 and 32 downregulated genes, respectively) compared to the use of all spots (15 upregulated genes; 16 downregulated genes) (Fig. 3e). PST can determine 81 genes compared to only 31 genes found by pseudotime method, and these gene sets are enriched for biological processes related to cancer (Fig. 3f). Overall, the incorporation of spatial information allowed us to capture spatial heterogeneity which revealed more significant molecular mechanisms within an individual tumour by reconstructing the evolutionary trajectories based on gene expression integrated with spatial information.

### 5. stLearn spatial cell-cell interactions

To study cell-cell interactions (CCI), we developed a method to combine cell type diversity and L-R co-expression into an interaction measure that can be used to automatically scan a whole tissue section. We computed L-R expression and cell type diversity independently (Fig. 1d), and then combined these two analysis streams at the end. First, we applied CellPhoneDB^29^ to the SMEclust results, to find L-R pairs that significantly contributes to cell-cell interactions. We scored spatial L-R co-expression in every spot, and then cluster spots based on L-R interactions (Fig. 4b, 4c). Second, cell type diversity was calculated based on the number of cell types present in a unit area of the tissue (Fig. 4a, 4d, 4e). Finally, by combining cell type diversity and L-R expression information, we identified regions with the highest likelihood of having the interactions (i.e. those with high L-R interaction and diverse cell types) (Fig. 4f)^12, 48, 49^. Details of the algorithms used are described in the methods section.

**Figure 4.**
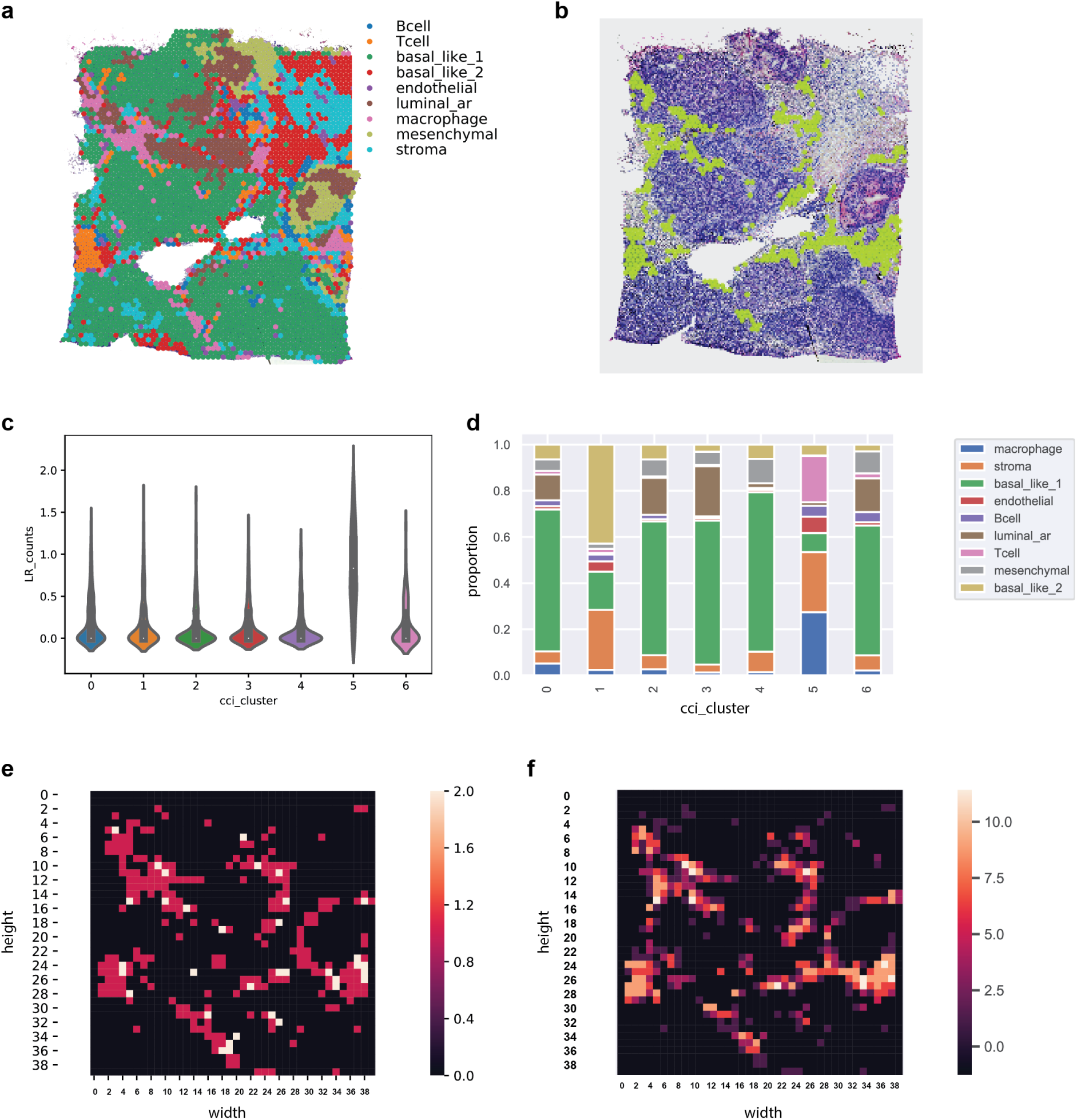
Inferring cancer-immune cell interactions by ligand-receptor (L-R) interactions in breast cancer. **a**, Spatial spots were assigned to different cell types via label transfer^18^ from a published breast cancer dataset^47^, which identified immune cells (B cell, T cell, macrophage) and non-immune cells within the tissue area; **b**, Seven spot clusters differing in their intensity of the neighbour L-R activity were detected. For each spot, the number of neighbouring spots that expressed a ligand or a receptor in each significant L-R pair was counted. The count data was combined to form a cell-cell interaction (CCI) matrix, which represented neighbourhood co-expression of each of the significant L-R pairs for all spots in the tissue. Of these seven clusters, cluster five was found to have the highest significant L-R expression (shown in green) **c**, An exemplar interaction analysis for a significant L-R pair (*CXCR*4*CXCL*12), with the co-expression activities spread across the seven clusters. The number of L-R neighbours for (*CXCR*4*CXCL*12) is shown, with the highest co-expression value found in cluster 5. **d**, Cell type distribution among the seven CCI clusters. Cluster 5 has a large accumulation of immune cell types. **e**, Cell type diversity distribution of three immune cell types (B cells, T cells, and macrophages) is shown as a heatmap. The heatmap scale shows the diversity score, that is, the number of different cell types in each unit area of the tissue. The immune cell type spatial density indicates potential immune-interacting regions across the whole tissue. **f**, L-R cluster with highest L-R activities (as shown in b) and cell type heterogeneity (as shown in e) were merged to define hotspots in the tissue where there was a diverse mix of immune cells and high L-R co-expression activities, suggesting regions with high cell-cell communications.

#### Case study 4

Here we worked with the same breast cancer dataset used for case study 3. This breast cancer tissue was staged as group II cancer with DCIS, IDC, and invasive ductal carcinoma, and lobular carcinoma *in situ* (LBIS). We used label transfer to annotate spots in our dataset based on another published cancer dataset^47^ (Fig. 4a), and compared these cell type annotations to the results of the CCI analysis. We found seven CCI clusters. Of these, CCI cluster 5 had the most predicted interactions, and displayed a unique L-R pattern compared to other clusters (Fig. 4c). When comparing cell type distributions within each cluster, we found that cluster 5 best aligned with the immune cells. A stacked bar plot of cell type composition across all CCI clusters (Fig. 4d) showed that cluster 5 also contained more immune cells (B cells, T cells, and macrophages) than other clusters. Regions with higher cell type diversity were determined using a window approach. Cell type diversity was computed as the number of cell types in a fixed size window (Fig. 1d, 4e). By integrating CCI clustering with cell type diversity, we identified regions with both high L-R interaction and more diverse cell types. These regions with high cell diversity and L-R activities are considered as hotspots in the tissue with most likely cell-cell interaction activities (Fig. 4f)^12, 48, 49^. To investigate biological inferences from the CCI analysis, we checked the co-expression pattern of a series of gene pairs, including known tumour-immune interacting pairs and control pairs, not involved in tumour-immune interactions. We found known cancer-related L-R pairs showed similar patterns to our CCI hotspot results (Supplementary Fig. 10), while the two control pairs did not show a spatially defined pattern (Fig. S10e, f).

### 6. Other functionalities in stLearn – Microenvironment detection

stLearn implements matrix factorisation methods to delineate a tissue section into distinct microenvironments. Microenvironment detection is implemented in stLearn via factor analysis of gene expression data. The user is then presented with the factors computed by stLearn, and can manually compare these factor loadings (weighted gene expression values in a latent dimension) with the known tissue morphology (Supplementary Fig. 11). Specifically, stLearn implements the following factor analysis options: PCA, ICA, and FA to find latent data dimensions (factors) that are correlated with biological or technical variation. We implemented an interactive visualisation function in stLearn to allow viewing of factor loadings across spots within the tissues. Where certain regions in the tissue show spatial patterns of factor loading levels, it can be inferred that these correspond to a microenvironment representing an anatomical region or a biological process. We also introduced a microenvironment detection analysis to find genes contributing the most to a factor of interest by finding expression values that are highly correlated with the factor loadings. With this approach, we identified genes drivers that contributed the most to defining the microenvironment.

#### Case study 5

Using a mouse traumatic brain injury model as described in our recent publication^50^, we aimed to find microenvironments that represent responses to tissue damage and anatomical regions that correspond to brain morphological structure. Spatial data was generated by the Legacy Spatial Transcriptomics platform, which measured fewer and bigger spots^16^ (also supported by stLearn). We compared two murine brain samples within a single multiplexed ST array; both samples contained injured and non-injured regions and one tissue had received a drug treatment prior to sampling. First, we applied factor analysis to the gene expression count matrix to project the original data into a latent space with 20 dimensions (factors). We used a interactive visualisation function in stLearn to inspect factor loadings to detect spatial patterns indicative of different regions within the brain tissue. In this step, we focused on two microenvironments for which consistent alignment of tissue morphology with with factor loadings were found. These include a hippocampus region (factor 3) and a damaged region (factor 1) (Supplementary Fig. 11a). In the second step, we used the Spearman’s rank correlation to find significant genes that correlate most with the identified factors. Next, these genes are likely drivers that determine the spatial pattern of the given microenvironment. The significant genes were analysed for functional enrichment using Enrichr with the Mouse Gene Atlas database. The enrichment result showed the hippocampus regions matched the activity map of factor 3, while the presence of immune cells such as microglia and macrophage cells matched the activity map of factor 1 (Supplementary Fig. 11b). The functional enrichment for these correlated genes was consistent with the known immune responses to damage^50^. Here as a proof of concept of the stLearn’s microenvironment detection algorithm, we completed the factor analysis method to identify an activity map with spatial gene expression within the tissue. We showed that the identified genes were enriched for biological pathways relevant to the morphology or treatment conditions of the tissue (Supplementary Fig. 11b).

## Discussion

### Normalisation and Clustering

To our knowledge, this study is the first to use information derived from tissue morphology to normalise gene expression values and improve downstream analysis of tissue biology. Our strategy is based on the main assumption supported by previous molecular studies. The assumption is that cells with more similar morphological features (e.g. cell size, nuclei size, granularity, or distribution) also have more similar transcriptional profiles^23^. We devised a creative method that utilises transfer learning with a pre-trained convolutional neural network to compute morphological similarities to improve the detection sensitivity of small clusters (Fig. 2, Supplementary Figs. 1-4). Previously, we developed neural network models that combined imaging pixel information and gene expression. The combinatorial approach outperformed the use of either gene expression or the tissue image alone^24^. Here, we use neighbourhood information to build a model that can denoise the technical variation inherent to single-cell or spatial sequencing, where lowly expressed genes are not detected or are highly variable. This approach has been suggested as a potential way to find spatial gene expression patterns^8^. Our novel strategy could adjust gene expression values to better reflect the cell types as shown by the improvement in detecting cell types and annatomical regions in mouse brain and human brain tissues.

By using SME normalised data, SMEclust incorporates the spatial dimensionality, tissue morphology and transcriptome-wide gene expression profiles to find cell types. Currently, scRNAseq is the method of choice for high resolution transcriptomic profiling of cell types and cell states in tissue. SMEclust surpasses scRNAseq clustering in several key features. The choice of clustering resolution for finding sub-clusters is still selected arbitrarily in most clustering methods using scRNAseq data^51^. In stLearn, we made unambiguous decision for sub-clustering by separating sub-clusters by their spatial distribution within a tissue. For example, in the breast cancer case study 3, only a single ductal cancer cluster was identified when using gene expression alone. However, the spatial separation of the cells within this large cluster provided the supporting evidence to justify splitting the cluster into three sub-clusters with transcriptionally different states and which followed different PST trajectories (Fig. 3). Work from single-molecule RNA insitu hybridyzation (osmFISH) shows that spatial colocalisation enabled the identification of rare cell-types that could not be detected by scRNASeq^52^. We showed SMEclust could identify rare cell types in the mouse brain tissues and at least 2-3 rare cell types were not detected by other methods that only used expression data, excluding spatial or morphological information.

### Pseudo-space-time (PST)

Intratumour heterogeneity (ITH) is a significant problem in oncology and can lead to low treatment effectiveness and high tumour recurrence. scRNASeq has proved a useful technology for the study of ITH^53, 54^ and reconstruction of the evolutionary^5^ trajectories of cancer cell types, making it a promising strategy to explore drug resistance in cancer therapy^55, 56^. However, this technology loses valuable data about the spatial location of cells, and thus cannot spatio-temporally track the evolutionary progress of tumour cell populations exhibiting ITH^8^.Efforts to spatially characterise the heterogeneity of subclonal cancers by multi-region sequencing exploited spatial patterns^57^ from a single time point within a tumour tissue^58, 59^. These studies observed convergent tumour evolution, characterised by branched evolutionary growth and multiple recurring, yet distinctive, inactivating mutations of the same tumour suppressor genes in different branches and regions^4^. Nevertheless, essential questions about tumour dynamics and evolution have remain elusive^8^, and the limitations of pseudotime analysis still exist^60^. To address these challenges, we developed pseudo-space-time method (Fig. 3b) which integrates spatial location with gene expression profiles to determine the spatial structure of subclone-specific evolution within (local level) and between (global level) tumour cell types/states. PST was inspired by the concept of spatiotemporal reconstruction of tumour ontogeny^39^ and pseudotime^27^ and diffusion-time^36^ analyses (Supplementary Fig. 5). This PST algorithm provides valuable information about branching evolution in cancer progression, and can be readily applied to ST data from developmental biology or related fields where cell state progression is expected.

### Cell-cell interactions

CCI analysis is challenging yet key for understanding complex tissue ecosystem dynamics. To address this challenge analytical methods like SpaOTsc^7^ and CellphoneDB^29^ have been developed. SpaOTsc^7^ uses an optimal transport algorithm to derive communications between signal senders and receivers. CellPhoneDB^29^ builds a curated L-R database to facilitate comparisons of these L-R interactions among clusters to find the most significant interacting pairs within the dataset. However, until now there has been no available method that combined both spatial cell-type distribution and L-R interactions to find hotspots within a tissue that likely have high CCI activities. stLearn CCI automatically scans for regions with high cell type densities that also show L-R co-expression; the co-occurrence of these two traits is suggestive of a highly interactive region^12, 48, 49^. The CCI algorithm combines multiple sources of information, including a database of known L-R pairs, gene expression, spatial location, and spatial cell-type distribution. stLearn’s CCI algorithm can find L-R pairs and spatial regions which are highly biologically relevant, like the cancer-immune cell interactions as in our breast cancer case study.

We have demonstrated stLearn’s three major algorithms with ST data generated from human and mouse, using both Visium and Legacy Spatial Transcriptomics data. Our stLearn analytical package is applicable with any ST data from any species or biological system, as long as tissue morphology, spatial location, and gene expression are simultaneously captured. Without tissue morphology information in HE image, stLearn is still useful to integrate spatial location and gene expression data, such as those from slide-seq^14^ and MERFISH^13^. The algorithms described in this work are pioneering analytical methods to help researchers study cells in morphologically intact tissue, and will be and can be readily applied to answer major questions about diseases and general biology.

## Methods

### 1. Spatial normalisation

ST provides transcriptome-wide gene expression profile with additional spatial location information and tissue morphology in HE tissue images. We developed this novel SME normalisation method to incorporate these two additional data types to adjust gene expression values between spots within a tissue. We used image processing functionalities and a neural network model for this analysis.

#### Spatial location

We implemented a disk smoothing method to utilise spatial location information to select neighbouring spot pairs for normalisation. For a given spot *S*_*i*_, a spot *S* _*j*_ is considered to be a neighbour if the centre-to-centre physical distance between the two spots *PD*_*i j*_ is shorter than a specified radial distance *r* as shown in equation (1). All paired spots *S*_*i*_ and *S* _*j*_ are then included as input to adjust for the gene expression of the centre spot *S*_*i*_ as described below.

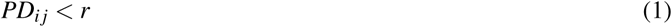

#### Morphological similarity

We measured the morphological similarity of paired spots *S*_*i*_ and *S* _*j*_ by the distance calculated based on two numeric vectors extracted from their corresponding spot images, i.e. the H&E tile that covers these spots. The high-level features of these spot images are extracted by ResNet50^61^, which is a well-established convolutional neural network (CNN) model widely used for image classification in computer vision. We used a transfer learning strategy, where the weights in ResNet50 were pre-trained using the ImageNet^62^ dataset (millions of images) and the final classification layer was extracted. This method leverages the generalisability of the pre-trained model in a large imaging dataset to extract numeric features from a new image^63^. As a result, the model can convert an image into a 2048-dimensional latent vector. We further applied PCA to extract the first 50 PCs as latent features to represent the spot morphology. The morphological distance *MD* of a centre spot *S*_*i*_ and its neighbour spot *S* _*j*_ can be calculated by cosine distance (other options such as Euclidean or Pearson distances are also available in stLearn), computed as in equation (2):

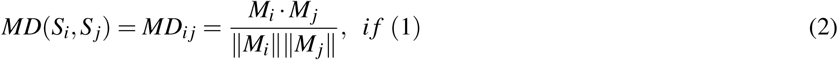

where *M*_*i*_ and *M*_*j*_ represent the morphological latent features for spots *S*_*i*_ and *S* _*j*_.

Incorporating both spatial location and morphological similarity, SME normalises gene expression of each spot *S*_*i*_ by:

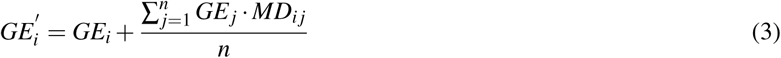

where 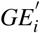 represents SME normalised gene expression for a centre spot *S*_*i*_. *GE*_*i*_ and *GE*_*j*_ represent raw gene expression for spot *S*_*i*_ and its *n* neighbour spots *S* _*j*_.

### 2. Spatial clustering algorithms – SMEclust

scRNAseq data has led to the development of numerous cell type identification methods using data-driven clustering^51^. These clustering methods, such those in the popular scanpy^31^ and Seurat^18^ pipelines, only use the gene expression information. stLearn clustering implements a novel algorithm that can incorporate the morphology and physical distance between measured spots.

#### Global clustering

Two levels of clustering are calculated: global and local. The global clustering algorithm is to find all clusters as described below :

1. Preprocess raw expression data by removing low abundant genes (detected in less than 1% of all spots) and proceed to option 1 or option 2. Option 1: perform SME normalisation, move to step 2, skip step 3, and move to step 4, 5, and 6. Option 2: perform another normalisation methods, move to step 2, 3, 4, 5, and 6.
2. Perform dimensionality reduction with PCA/UMAP
3. Adjust the PCA/UMAP expression space by using the disk smoothing approach, where the spatial information of proximal spots is used. For each spot, stLearn scans neighbouring spots that are within a specified distance (using cKFTree query). The PCA/UMAP coordinates of the spots are then adjusted based on the identified neighbours by calculating the mean values of the coordinates adjusted by morphological distance (*MD*_*i j*_ as in formula 3): *ad justed*_*coordinates* = (*neighbouring*_*coordinates* \**MD*_*i j*_)*/*(*number o f neighbours*)
4. Construct kNN graph based on Euclidean distance
5. Run Louvain or K-means clustering
6. Run local clustering to find sub-clusters that are spatially separated and to merge spots that are spatially collocated as described below.

#### Local clustering

At local scale, we use spatial location information of spots within a tissue section to refine the global clustering result by sub-clustering and by merging singleton spots or small clusters. Sub-clustering is performed when one cluster consists of spots located at different parts of the tissue. Merging spots is to assign a local clustering label for spots that do not belong to any global cluster. The assumption for local clustering is that spots of a cell type or sub-type that are geographical neighbours in the tissue will be more similar than those that are physically distant.

Details of local clustering steps for ST data is described below:

1. Using spatial coordinates, for every spot, find neibouring spots in the *ε* distance within the tissue, and identify location-based core spots with more than *min*_*samples* neighbours. We used DBSCAN to find location-based clusters of arbitrarily shape even in the presence of outliers. We recommend that *ε* = 100 for high resolution images from Visium data, but *ε* should be varied depending on the sequencing platform used, due to the size of the image. Specific recommendations are provided in the stLearn documentation.
2. For each global cluster, identify the location-based clusters belonging to the global cluster. If a global cluster correspond to more than one location-based clusters, the global cluster can be split into sub-clusters.
3. Assign each location-based non-core spot to a nearby location-based cluster if the cluster has at least *ε* neighbours, if not then set the spot to be a singleton without a clustering label.

### 3. Tissue trajectory analysis by pseudo-space-time algorithms

#### Pseudo-space-time (PST) analysis algorithms

We developed a novel algorithm to calculate PSTD between between two sets of slide spots to model gradient changes in transcriptional states. In this section, we will describe two levels of spatial trajectory inferences based on PSTD. The first is the global level analysis which explores the relationship between two clusters. Secondly, the local level can be used to reveal the relationship between sub-clusters of a single cluster.

#### Calculating pseudo-space-time distance

Given two sets of spots *U* and *V* with their corresponding centroids *u* and *v*, we applied the following formulae:

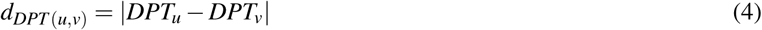

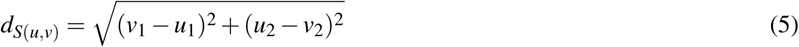

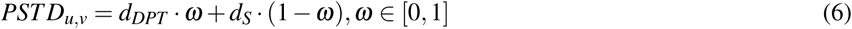

where (*u*_1_, *u*_2_) and (*v*_1_, *v*_2_) are the coordinates of the centroids of clusters *U* and *V*; *DPT*_*u*_ and *DPT*_*v*_ are the maximum DPT scores among spots in each cluster, and *ω* is a weighting factor reflecting the balance between gene expression and physical distance.

The maximum DPT score for all spots in each cluster is used as a representative value for the cluster because it reflects the difference in gene expression. We found that, even at the sub-cluster level, where spots are most similar, there still exists a dynamic variation of gene expression among all spots. For example, the distribution of DPT values in different sub-clusters is stochastic (Supplementary Fig. 7c). Therefore, the single mean value or the summation of all individual spots is not sufficient to represent the transcriptional states of sub-clusters, and the maximum value therefore appears to be a better measure. A reasonable assumption for transcriptional states of individual spots within a sub-cluster is that two spots that have a greater physical distance within a sub-cluster are more transcriptionally different than two more proximal spots. Then, *d*_*s*_ reflects the relationship of two sub-clusters by calculating the distance between them. Another parameter is *ω* which represents the weight with which gene expression effect and the spatial distance effect contribute to calculate the *PSTD*. When *ω*=0, the DPT distance is identical to physical distance and does not take into account gene expression distance. When *ω*=1, the spatial distances will not be taken into account and the graph will be optimised based on the DPT distances only. Intermediate values of *ω* allow the user to adjust the relative contributions of these two measures to graph optimisation.

Applying the formulae above, we built a PSTD matrix for input to the PST analysis method at the local and global levels.

#### Local trajectory inference

At the local level, a selected cluster (*M*) may be divided into multiple sub-clusters (*m*_1_, *m*_2_, *m*_3_, *…, m*_*n*_) if it occupies more than one physical location. The PSTD matrix can be used to determine the weight of the trajectory line connecting each pair of sub-clusters. The line illustrates the spatial-transcriptional relationship between the pair of clusters. The trajectory directionality can be assigned using the DPT distance. If *DPTdistance* > 0 then we assign forward direction, if *DPTdistance <* 0 then we assign reverse direction. Therefore, by using the PSTD and *DPT*, we can infer the trajectory from a sub-cluster (as root) to other sub-clusters within the cluster *M*.

#### Global trajectory inference

While local trajectory analysis can define biological processes within one cluster, it does not allow for comparison between clusters. We therefore developed the second part of the PST algorithm to find how multiple clusters are connected within a tissue (Fig. 1b). Given two sets of sub-clusters: *u*_1_, *u*_2_, *u*_3_, *…, u*_*n*_ and *v*_1_, *v*_2_, *v*_3_, *…, v*_*n*_ in two separate clusters *U* and *V*, the global PST is described below:

1. At the cluster level, we can first order sub-clusters in each of these two clusters by ranking the sub-cluster’s average spot DPT values (Fig. 4a).
2. On the assumption that the overall DPT order is from *U* to *V*, we denote that the tissue evolution starts from *U* to *V*.
3. We built the PSTD matrix with dimension as the number of *u*_*i*_ nodes in *U* and *v* _*j*_ nodes in *V* from two sets of sub-clusters *u*_1_, *u*_2_, *u*_3_, *… u*_*n*_ and *v*_1_, *v*_2_, *v*_3_, *… v*_*n*_. This means that the values of the distance matrix are a set of PSTD between every two sub-clusters *u*_*i*_ and *v* _*j*_. We compressed each sub-cluster to become a node in the graph and the distance between two nodes (*u*_*n*_ -> *v*_*n*_) is PSTD computed as below. The *d*_*DPT*_ is based on the subtraction of the average of DPT scores for all spots in each of the two nodes. The *d*_*s*_ is the physical distance between centroids of spot coordinates in *u*_*n*_ and *v*_*n*_.
4. From the *PSTD* matrix, we built a fully connected, directed graph comprising of the initial trajectories that capture the directions from sub-clusters of *U* to sub-clusters of *V*. For example, a fully connected graph is constructed with *D*: (*u*_1_, *v*_1_), (*u*_1_, *v*_2_), (*u*_1_, *v*_3_), *…* and (*u*_2_, *v*_1_), (*u*_2_, *v*_2_), (*u*_2_, *v*_3_), *…*, (*u*_*n*_, *v*_*n*_).
5. From the fully connected directed graph, we found an optimal branching structure of the graph to construct the evolutionary trajectories. A pseudo-root is added to the graph to form an arborescence (a rooted, directed tree), which can be optimised by using a minimum directed spanning tree approach with Chu–Liu/Edmonds’ algorithm^37^. As the default, we set the pseudo-root *r* as the centroids of all sub-clusters in the first layer. The minimum directed spanning tree approach is described in the later section,
6. After finding the minimum directed spanning tree, we obtained the optimal graph, for example, *D*: (*u*_1_, *v*_1_), (*u*_2_, *v*_2_), (*u*_2_, *v*_3_), which represents the trajectories of each sub-cluster from the lower layer to the higher layer. With this approach, one node can be the start node of multiple branches but be the end node of one branch only. Finally, we plotted the branches to the tissue image to allow for the visualisation of the evolutionary trajectories.

We applied optimum branching (with Chu–Liu/Edmonds’ algorithm^37^) as a directed analogue of the minimum spanning tree problem to the PST scenario. This yields a weighted, directed graph *D*(*V, E*) where *V* is the set of nodes (sub-clusters), *E* is the set of directed edges (raw trajectories), a node *r* called root (assigned pseudo-root) in *V*, and *ω* is the weight of each edge in *E* (calculated as PSTD). From the ‘raw’ fully connected tree, we identified a directed spanning tree or spanning arborescence *A* with a root at *r* such that every node in *A* has two edges (except for the tip of the branch, which has one edge). The optimisation process is performed such that *A* has a minimum weight, defined as the sum of all *ω* in *A* as the cost function:

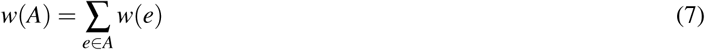

By letting *F* be a set of *n* − 1 edges extracted from *D* by determining the cheapest edge (an edge with the lowest weight) entering each node *v* ≠ *r* that always forms a path *v*_0_ ⟵ *v*_1_ ⟵*…* ⟵ *v*_*n*_, where each *v*_*i*_ is an original node. Because of the setup of the graph, we obtained *F* as a directed acyclic graph the simplest scenario (without any cycle in the graph) of this algorithm. In detail, for the initialisation step, *F* is empty. At each step, the algorithm selects an arbitrary node *v* ≠ *r*, which does not yet have an incoming edge in *F*, finds the cheapest edge (*u, v*) *∈E* entering *v*, and adds (*u, v*) to *F*. To find the cheapest edge entering a given node, the algorithm repeatedly executes minimum weighted edge extraction operations until the returned edge is not a self-loop in the current graph.

#### Trajectory-based transition gene detection

As a part of the PST analysis, stLearn allows users to find genes that likely drive the transcriptional trajectories. For a chosen clade *B*_*i*_ and for a set of genes *E* with gene expression values *e*_1_, *e*_2_, *…e*_*n*_, genes that are differentially expressed between the start and end nodes in a trajectory pathway are tested for significance using Spearman’s rank correlation between gene expression value and pseudotime. Spearman’s rank correlation coefficient for *r* _*j*_ is high if the gene expression *e* _*j*_ is linearly correlated with *DPT*_*j*_ in clade *B*_*i*_. After correcting for multiple testing, significant genes that are identified are likely to be trajectory driving genes. Genes with Spearman’s rank correlation coefficient higher than a specified threshold (0.2 by default) are determined as upregulated transition genes and vice versa. In contrast, genes with correlation coefficient lower than the threshold are identified as downregulated transition genes.

#### Gene set enrichment analyses

We performed functional enrichment analyses for the top 100 upregulated and downregulated transition genes which were identified by trajectory-based differential expression analysis. All three Gene Ontology (GO) categories were used: molecular function, biological process, and cellular component. This step was implemented using gseapy^64^ and Enrichr^65^.

### 4. Cell-cell interaction algorithms

Cell-cell interaction (CCI) analysis consists of two main steps that allow us to incorporate spatial information in the search for tissue regions with high CCI activities.

First, CCI finds cell type diversity across the tissue. Each spot in the tissue sample can be annotated by label transfer from any related publicly available dataset using Seurat v3^18^. The tissue space is divided into small windows based on user-specified number of divisions along each edge of the tissue section. To calculate the density of cell types of interest (for example immune cell types), we annotate cell types in every spatial spot, and count the number of cell types for all spots within in each window. The count represents the cell type density over a fixed area of tissue, thus representing cell type diversity. Cell type density can be shown in a table or as a heatmap as as outlined in our online tutorial.

Second, CCI finds L-R co-expression between neighbouring spots. Permutation tests for the enrichment of L-R pairs between two cell types compared to the random null distribution are first performed using CellPhoneDB^29^. Significant L-R pairs are then used for calculating cell-cell interactions. For each spot, the nearest neighbours within a specified distance are scanned with cKDTree, in order to check whether they express ligand or receptor genes above a specified threshold. Thus, reliable expression detection among neighbours is ensured. The proportion of neighbouring spots co-expressing L-R pairs for the central spot are then calculated as in equation (8) and used for clustering analysis.

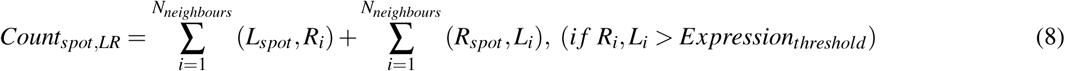

where for a spot containing a positive ligand expression, the (*L*_*spot*_, *R*_*i*_) is the count of neighbouring spots with the receptor expression value higher than an expression *threshold*, and similarly for (*R*_*spot*_, *L*_*i*_).

We can then form a CCI matrix, containing all significant L-R pairs as columns and spot coordinates as rows. This matrix allows us to cluster tissue regions by grouping the spots with the most similar L-R co-expression values. Finally, we combine CCI measures with cell type diversity measures to identify regions that are both the most diverse and have high L-R co-expression, indicating tissue areas that are most likely to harbour active cell-cell interactions.

We use a scanning window approach to combine the findings from cell type heterogeneity and from L-R co-expression between neighbouring cells. The aim is to find highly interacting spatial regions with high diversity and high L-R co-expression. Cell type heterogeneity scores in each window are transformed to z-scores. The window approach is applied to the LR co-expression clusters, by counting the percentage of spots belonging to the largest expressed CCI cluster in each window. The counts are then also transformed into z-scores, and the mean of the two z-scores in each window provides the final score that can be used to find locations with the top combined scores, which can be plotted as a heatmap for the whole tissue.

### 5. Details of the spatial trajectory analysis of breast cancer tissue

For the breast cancer ST data, we performed SME normalisation and SMEclust as described above. To annotate clusters, we identified differentially expressed marker genes and compared these markers to those present in the auxiliary breast cancer scRNAseq dataset^47, 66^ (Supplementary Fig. 6b, 7a, 7b, 7c). We identified eleven broad clusters, seven of which are predicted to contain invasive cancer cells. Some of these clusters were distributed across different locations within the tissue, so we resolved these into spatially-distributed sub-clusters with SMEclust as described above. Of particular interest were the DCIS cluster (marked as pink in Supplementary Fig. 6b) which split into three morphologically-distinct sub-clusters (clusters 6, 11 and 14 in Supplementary Fig. 6d) and the IDC cluster (marked as sky blue in Supplementary Fig. 6b) which split into seven sub-clusters and likely comprises invasive cancer and immune/lymphoid clusters.

We then used trajectory analysis to study DCIS-IDC progression. As we expected DCIS to represent an earlier stage of cancer progression than IDC, we set DCIS as the trajectory root. In the PAGA graph showing the relationship among clusters based on gene expression, the unsegregated DCIS and IDC clusters share a connected edge in the graph and are close to each other in dimensionality reduction space (Supplementary Fig. 7d). We inferred pseudotime distance for each cluster relative to the DCIS root; the gradient of DPT scores is shown in the PAGA embedding space (Supplementary Fig. 7e) and spatial coordinate (Supplementary Fig. 6c). As predicted, this gradient suggested the likely transition of DCIS-IDC in the tissue.

As the common ancestor cell that gave rise to the DCIS-IDC progression is unknown, we tried to infer the subclonal cancer evolution based on the combination of DPT and physical distance information to generated a PSTD matrix as described above. We constructed trajectories for each of the three DCIS sub-clusters (Supplementary Fig. 6e) with the pairwise sub-clusters connections and distances: 11-6 (0.813), 11-14 (0.347), 14 – 6 (0.833) corresponding to how far the relationship is. Trajectory direction was defined by the distance between the maximum DPT of each sub-cluster.

### 6. Implementation of software

Here we describe the technical detail for the stLearn software, which incorporates three key, novel analysis algorithms. stLearn software is publicly available at PyPI and the software documentation can be found at https://stlearn.readthedocs.io/.

stLearn’s core is constructed based on tensorflow^67^ and Pillow by Alex Clark for deep learning and image processing modules; scipy^68^ for the spatial analysis module; scanpy^31^ for the gene expression analyses modules; networkx^69^ and matplotlib^70^ for network analysis and visualisation; and anndata^31^ for the main object to store and process data. MATLAB software Histology (GitHub: https://github.com/petersaj/histology) is used for comparing the SMEclust results to an anatomical reference. As stLearn is a Python-based software, it is compatible with machine learning or deep learning packages like scikit-learn^71^, tensorflow. Users can also can utilise functions from popular image processing packages, such as OpenCV^72^ or Pillow for further analysis of tissue images. stLearn uses annData object, compatible to many other R/Python software which uses the common SingleCellExperiment object.

## Acknowledgements

We thank members in Nguyen’s Genomics and Machine Learning Lab for helpful discussion. This work has been supported by the Australian Research Council (ARC DECRA DE190100116), the University of Queensland, and the Genome Innovation Hub.

## Author contributions statement

Q.N., D.P., X.T., J.X. conceived the experiments and developed the algorithms. D.P., X.T., J.X. wrote the software. D.P., X.T., J.X., Q.N., P.Y.L., L.F.G., J.V. and M.J.R. conducted the experiments and analysed the data. A.R., J.V. and M.J.R. helped with the interpretation of brain anatomy and breast cancer. Q.N., D.P., J.X, L.F.G. wrote the manuscript. All authors have reviewed and approved the manuscript.

## References

1. Regev, A. et al. Science forum: The human cell atlas. eLife 6, e27041 (2017).

2. Scadden, D. T. Nice neighborhood: emerging concepts of the stem cell niche. Cell 157, 41–50 (2014).

3. Janiszewska, M. The microcosmos of intratumor heterogeneity: the space-time of cancer evolution. Oncogene 39, 2031–2039 (2020).

4. Swanton, C. Intratumor heterogeneity: evolution through space and time. Cancer research 72, 4875–4882 (2012).

5. Greaves, M. & Maley, C. C. Clonal evolution in cancer. Nature 481, 306 (2012).

6. Tanay, A. & Regev, A. Scaling single-cell genomics from phenomenology to mechanism. Nature 541, 331–338 (2017).

7. Cang, Z. & Nie, Q. Inferring spatial and signaling relationships between cells from single cell transcriptomic data. Nat Commun 11, 2084 (2020).

8. Lähnemann, D. et al. Eleven grand challenges in single-cell data science. Genome biology 21, 1–35 (2020).

9. Burgess, D. J. Spatial transcriptomics coming of age. Nat. Rev. Genet. 20, 317 (2019).

10. Maynard, K. R. et al. Transcriptome-scale spatial gene expression in the human dorsolateral prefrontal cortex. bioRxiv DOI: 10.1101/2020.02.28.969931 (2020).

11. Beechem, J. M. High-Plex spatially resolved RNA and protein detection using digital spatial profiling: A technology designed for immuno-oncology biomarker discovery and translational research. Methods Mol. Biol. 2055, 563–583 (2020).

12. Eng, C. L. et al. Transcriptome-scale super-resolved imaging in tissues by RNA seqFISH. Nature 568, 235–239 (2019).

13. Chen, K. H., Boettiger, A. N., Moffitt, J. R., Wang, S. & Zhuang, X. RNA imaging. Spatially resolved, highly multiplexed RNA profiling in single cells. Science 348, aaa6090 (2015).

14. Rodriques, S. G. et al. Slide-seq: A scalable technology for measuring genome-wide expression at high spatial resolution. Science 363, 1463–1467 (2019).

15. Fan, Z., Chen, R. & Chen, X. SpatialDB: a database for spatially resolved transcriptomes. Nucleic Acids Res. 48, D233–D237 (2019).

16. Stahl, P. L. et al. Visualization and analysis of gene expression in tissue sections by spatial transcriptomics. Science 353, 78–82 (2016).

17. Queen, R., Cheung, K., Lisgo, S., Coxhead, J. & Cockell, S. Spaniel: analysis and interactive sharing of spatial transcriptomics data. bioRxiv DOI: 10.1101/619197 (2019).

18. Stuart, T. et al. Comprehensive integration of single-cell data. Cell 177, 1888–1902 (2019).

19. J, B. SpatialCPie: Cluster analysis of Spatial Transcriptomics data (2020). R package version 1.2.0.

20. Edsgard, D., Johnsson, P. & Sandberg, R. Identification of spatial expression trends in single-cell gene expression data. Nat. Methods 15, 339–342 (2018).

21. Svensson, V., Teichmann, S. A. & Stegle, O. SpatialDE: identification of spatially variable genes. Nat. Methods 15, 343–346 (2018).

22. Dries, R. et al. Giotto, a pipeline for integrative analysis and visualization of single-cell spatial transcriptomic data. bioRxiv DOI: 10.1101/701680 (2019).

23. Cutiongco, M. F. A., Jensen, B. S., Reynolds, P. M. & Gadegaard, N. Predicting gene expression using morphological cell responses to nanotopography. Nat Commun 11, 1384 (2020).

24. Tan, X., Su, A., Tran, M. & Nguyen, Q. SpaCell: integrating tissue morphology and spatial gene expression to predict disease cells. Bioinformatics 36, 2293–2294 (2020).

25. Saelens, W., Cannoodt, R., Todorov, H. & Saeys, Y. A comparison of single-cell trajectory inference methods. Nat. Biotechnol. 37, 547–554 (2019).

26. Waclaw, B. et al. A spatial model predicts that dispersal and cell turnover limit intratumour heterogeneity. Nature 525, 261–264 (2015).

27. Trapnell, C. et al. The dynamics and regulators of cell fate decisions are revealed by pseudotemporal ordering of single cells. Nat. biotechnology 32, 381 (2014).

28. Cabello-Aguilar, S. et al. SingleCellSignalR: inference of intercellular networks from single-cell transcriptomics. Nucleic Acids Res. (2020).

29. Efremova, M., Vento-Tormo, M., Teichmann, S. A. & Vento-Tormo, R. Cellphonedb: inferring cell–cell communication from combined expression of multi-subunit ligand–receptor complexes. Nat. Protoc. 15, 1484–1506 (2020).

30. He, K., Zhang, X., Ren, S. & Sun, J. Deep residual learning for image recognition. CoRR abs/1512.03385 (2015). 1512.03385.

31. Wolf, F. A., Angerer, P. & Theis, F. J. Scanpy: large-scale single-cell gene expression data analysis. Genome biology 19, 15 (2018).

32. Subelj, L. & Bajec, M. Unfolding communities in large complex networks: combining defensive and offensive label propagation for core extraction. Phys Rev E Stat Nonlin Soft Matter Phys 83, 036103 (2011).

33. Lein, E. S. et al. Genome-wide atlas of gene expression in the adult mouse brain. Nature 445, 168–176 (2007).

34. Shamash, P., Carandini, M., Harris, K. & Steinmetz, N. A tool for analyzing electrode tracks from slice histology. bioRxiv DOI: 10.1101/447995 (2018).

35. Wolf, F. A. et al. Paga: graph abstraction reconciles clustering with trajectory inference through a topology preserving map of single cells. Genome biology 20, 59 (2019).

36. Haghverdi, L., Buettner, M., Wolf, F. A., Buettner, F. & Theis, F. J. Diffusion pseudotime robustly reconstructs lineage branching. Nat. methods 13, 845 (2016).

37. Gabow, H. N., Galil, Z., Spencer, T. & Tarjan, R. E. Efficient algorithms for finding minimum spanning trees in undirected and directed graphs. Combinatorica 6, 109–122 (1986).

38. Gerlinger, M. et al. Intratumor heterogeneity and branched evolution revealed by multiregion sequencing. N Engl j Med 366, 883–892 (2012).

39. Sottoriva, A. et al. Intratumor heterogeneity in human glioblastoma reflects cancer evolutionary dynamics. Proc. Natl. Acad. Sci. 110, 4009–4014 (2013).

40. Walker, C., Mojares, E. & del Río Hernández, A. Role of extracellular matrix in development and cancer progression. Int. journal molecular sciences 19, 3028 (2018).

41. Han, C.-c. & Wan, F.-s. New insights into the role of endoplasmic reticulum stress in breast cancer metastasis. J. breast cancer 21, 354–362 (2018).

42. Belin, S. et al. Dysregulation of ribosome biogenesis and translational capacity is associated with tumor progression of human breast cancer cells. PloS one 4 (2009).

43. Noensie, E. N. & Dietz, H. C. A strategy for disease gene identification through nonsense-mediated mrna decay inhibition. Nat. biotechnology 19, 434–439 (2001).

44. Standish, L. J. et al. Breast cancer and the immune system. J. Soc. for Integr. Oncol. 6, 158 (2008).

45. Sistigu, A. et al. Cancer cell–autonomous contribution of type I interferon signaling to the efficacy of chemotherapy. Nat. medicine 20, 1301 (2014).

46. Budhwani, M., Mazzieri, R. & Dolcetti, R. Plasticity of type I interferon-mediated responses in cancer therapy: from anti-tumor immunity to resistance. Front. oncology 8, 322 (2018).

47. Karaayvaz, M. et al. Unravelling subclonal heterogeneity and aggressive disease states in TNBC through single-cell rna-seq. Nat. communications 9, 1–10 (2018).

48. Schapiro, D. et al. histoCAT: analysis of cell phenotypes and interactions in multiplex image cytometry data. Nat. Methods 14, 873–876 (2017).

49. Rieckmann, J. C. et al. Social network architecture of human immune cells unveiled by quantitative proteomics. Nat. Immunol. 18, 583–593 (2017).

50. Willis, E. F. et al. Repopulating microglia promote brain repair in an IL-6-dependent manner. Cell 180, 833–846 (2020).

51. Duo, A., Robinson, M. D. & Soneson, C. A systematic performance evaluation of clustering methods for single-cell RNA-seq data. F1000Res 7, 1141 (2018).

52. Codeluppi, S. et al. Spatial organization of the somatosensory cortex revealed by osmFISH. Nat. Methods 15, 932–935 (2018).

53. Shipitsin, M. et al. Molecular definition of breast tumor heterogeneity. Cancer cell 11, 259–273 (2007).

54. Marusyk, A. & Polyak, K. Tumor heterogeneity: causes and consequences. Biochimica et Biophys. Acta (BBA)-Reviews on Cancer 1805, 105–117 (2010).

55. Brady, S. W. et al. Combating subclonal evolution of resistant cancer phenotypes. Nat. communications 8, 1–15 (2017).

56. Lee, M. C. W. et al. Single-cell analyses of transcriptional heterogeneity during drug tolerance transition in cancer cells by rna sequencing. Proc. Natl. Acad. Sci. 111, E4726–E4735 (2014).

57. Casasent, A. K., Edgerton, M. & Navin, N. E. Genome evolution in ductal carcinoma in situ: invasion of the clones. The J. pathology 241, 208–218 (2017).

58. Navin, N. et al. Tumour evolution inferred by single-cell sequencing. Nature 472, 90 (2011).

59. Wang, Y. et al. Clonal evolution in breast cancer revealed by single nucleus genome sequencing. Nature 512, 155–160 (2014).

60. Wagner, D. E. & Klein, A. M. Lineage tracing meets single-cell omics: opportunities and challenges. Nat. Rev. Genet. 1–18 (2020).

61. He, K., Zhang, X., Ren, S. & Sun, J. Deep residual learning for image recognition. CoRR abs/1512.03385 (2015).

62. Deng, J. et al. ImageNet: A Large-Scale Hierarchical Image Database. In CVPR09 (2009).

63. Pan, S. J. & Yang, Q. A survey on transfer learning. IEEE Trans. on Knowl. Data Eng. 22, 1345–1359 (2010).

64. Subramanian, A. et al. Gene set enrichment analysis: A knowledge-based approach for interpreting genome-wide expression profiles. Proc. Natl. Acad. Sci. 102, 15545–15550 (2005).

65. Kuleshov, M. V. et al. Enrichr: a comprehensive gene set enrichment analysis web server 2016 update. Nucleic acids research 44, W90–W97 (2016).

66. Vickovic, S. et al. High-definition spatial transcriptomics for in situ tissue profiling. Nat. methods 16, 987–990 (2019).

67. Abadi, M. et al. Tensorflow: Large-scale machine learning on heterogeneous distributed systems. arXiv preprint arXiv:1603.04467 (2016).

68. Jones, E., Oliphant, T. & Peterson, P. Scipy: Open source scientific tools for python. (2001).

69. Hagberg, A., Swart, P. & S Chult, D. Exploring network structure, dynamics, and function using networkx. Tech. Rep., Los Alamos National Lab.(LANL), Los Alamos, NM (United States) (2008).

70. Hunter, J. D. Matplotlib: A 2D graphics environment. Comput. science & engineering 9, 90–95 (2007).

71. Pedregosa, F. et al. Scikit-learn: Machine learning in python. J. machine learning research 12, 2825–2830 (2011).

72. Bradski, G. & Kaehler, A. Learning OpenCV: Computer vision with the OpenCV library (“O’Reilly Media, Inc.”, 2008).

