## SupplementaryFigures for "stLearn: integrating spatial location, tissue morphology and gene expression to find cell types, cell-cell interactions and spatial trajectories within undissociated tissues"

### Supplementary Figures

*D. Pham, et al.*

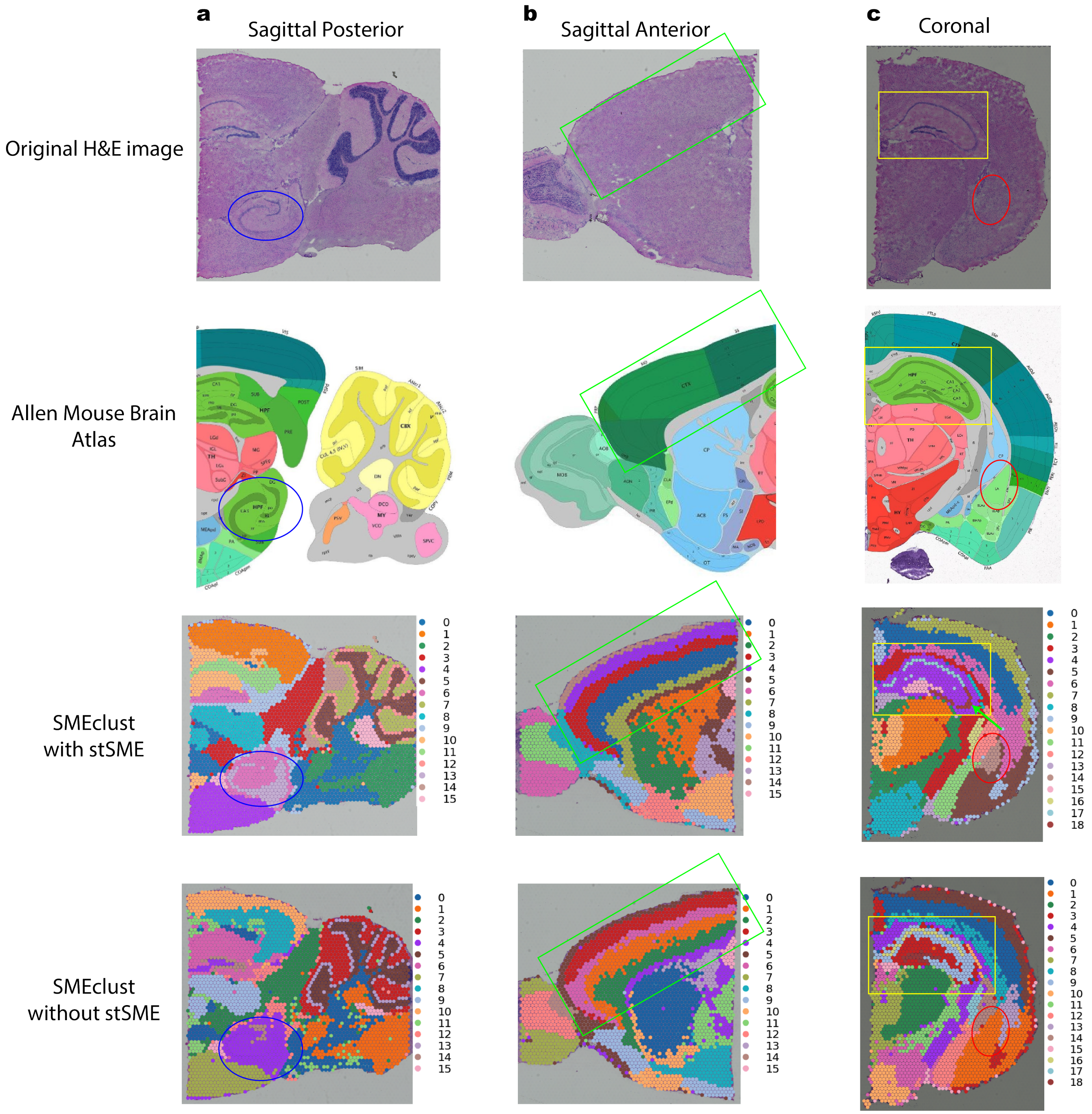


**Supplementary Figure 1.** Comparison of SMEclust performance with and without SME normalisation using 10x Visium mouse brain dataset. **a**, Sagittal posterior section; With SME normalisation, SMEclust can detect CA3sp (cornu ammonis 3, pyramidal layer)(blue circle) from the hippocampus. **b**, Sagittal anterior section; Clustering with SME normalisation shows clear and continuous cerebral cortex (CTX) layers (green box), which are less well-separated without SME normalisation. **c**, Coronal section; Comparison between clustering with and without SMEclust demonstrates the advantages of applying SMEclust in defining three main regions of the hippocampal region i.e. CA3, CA1/2 and dentate gyrus (DG). In addition, application of SMEclust also differentiated the lateral amygdala nucleaus (LA) from the cortical subplate (CTXsp) region (red circle) a lot more efficiently as compared to other methods. In all scenarios, SMEclust is the best data-driven clustering method that shows the highest consistency with the anatomical reference, and less noise.

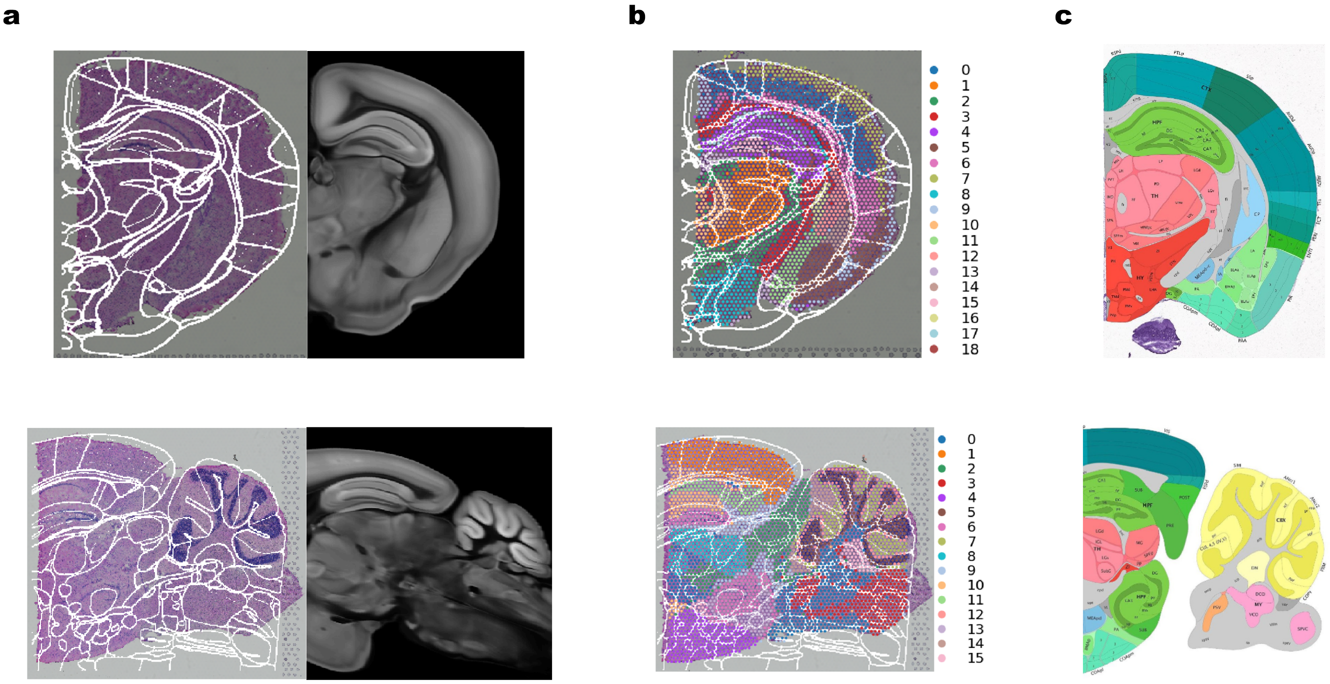


**Supplementary Figure 2.** Creating a customised, digital reference for anatomical regions within a mouse brain tissue section. **a**, A registration method based on a common coordinate framework (CCF). An image of a brain tissue section can be registered to a three-dimensional brain reference in the mouse brain CCF by deploying the Histology software (GitHub: `https://github.com/petersaj/histology`). Two examples with mouse brain coronal (top) and sagittal posterior (bottom) sections are shown. The two tissue sections were superimposed by anatomical structures (left) projected from the 3D Allen Mouse Brain Atlas reference (right). **b**, Mapping of the 3D Allen Mouse Brain Atlas to SMEclust results. The mapping results allowed for qualitative assessment of the expected anatomical regions that correspond to the clustering results. **c**, Generic and two dimensional references from the Allen Mouse Brain Atlas can also be used for comparing the clustering with expected anatomical regions.

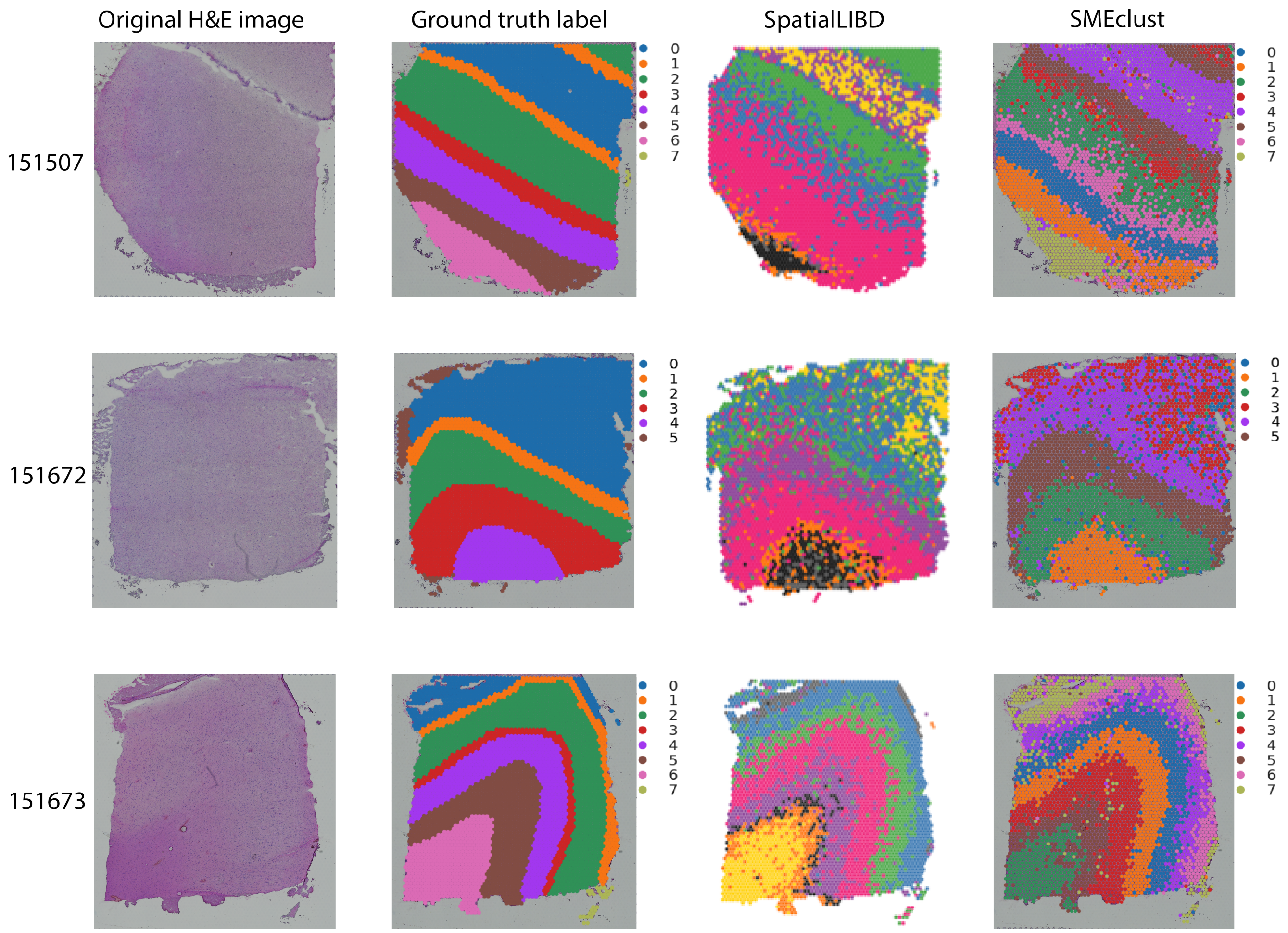


**Supplementary Figure 3.** Comparison of clustering performance of SpatialLIBD and SMEclust method using human dorsolateral prefrontal cortex Visium dataset[1].Three of the twelve tested datasets (IDs: 151507, 151672, 151673) are shown. In each row, from left to right, the H&E images, ground truth annotations, and clustering results using spatialLIBD and SMEclust are presented for comparisons. Ground truth annotations were manually assigned by a pathologist as reported in the original publication[1]. For all three samples shown, SMEclust results were closer to the ground truth than those generated by SpatialLIBD, showing less noise and detecting more cellular layers.

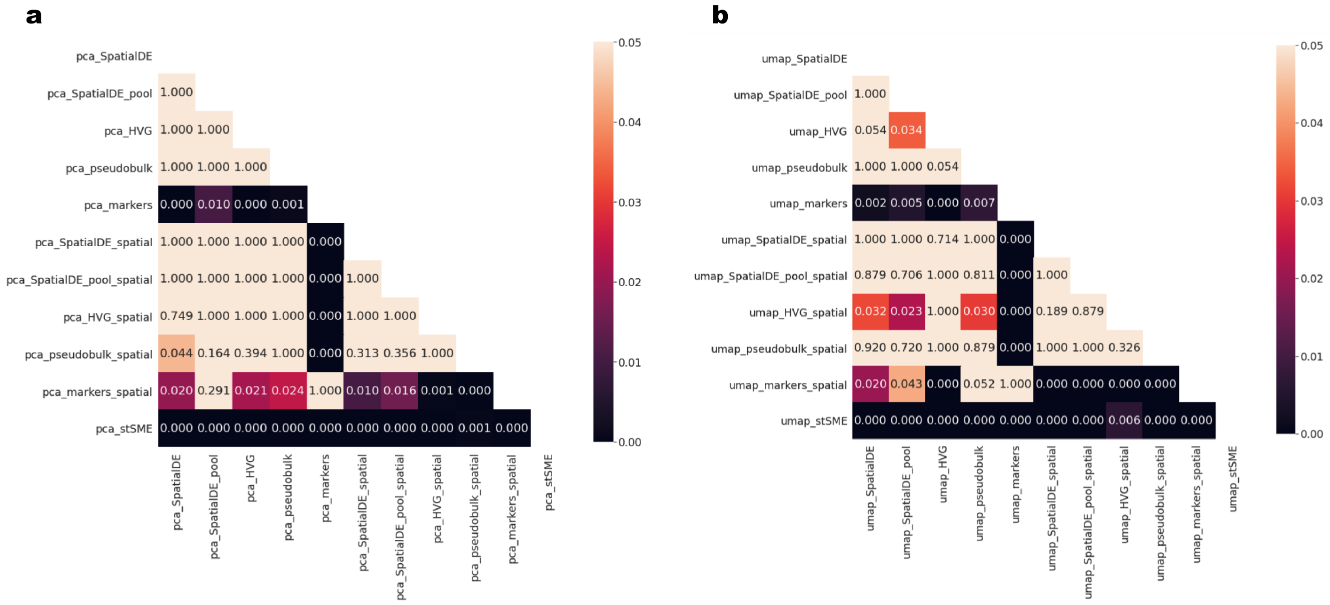


**Supplementary Figure 4.** Clustering comparison between SpatialLIBD and SMEclust method using 12 human dorsolateral prefrontal cortex Visium samples[1]. **a**, For each of the 12 samples, an Adjusted Rand Index (ARI) was calculated by comparing the clustering result from SpatialLIBD or SMEclust to the ground-truth clusters reported in the paper[1]. Ten data preprocessing options to select genes for clustering are shown in the x-axis and y-axis labels. The heatmap colours show P-values from F-tests for the differences in the distributions of ARI values between SpatialLIBD and SMEclust applied for the 12 tissue samples. The P-values were corrected for multiple testing. Panel **a** shows clustering methods applied on PCA space. **b**, similar to **a**, but clustering methods were applied on UMAP space.

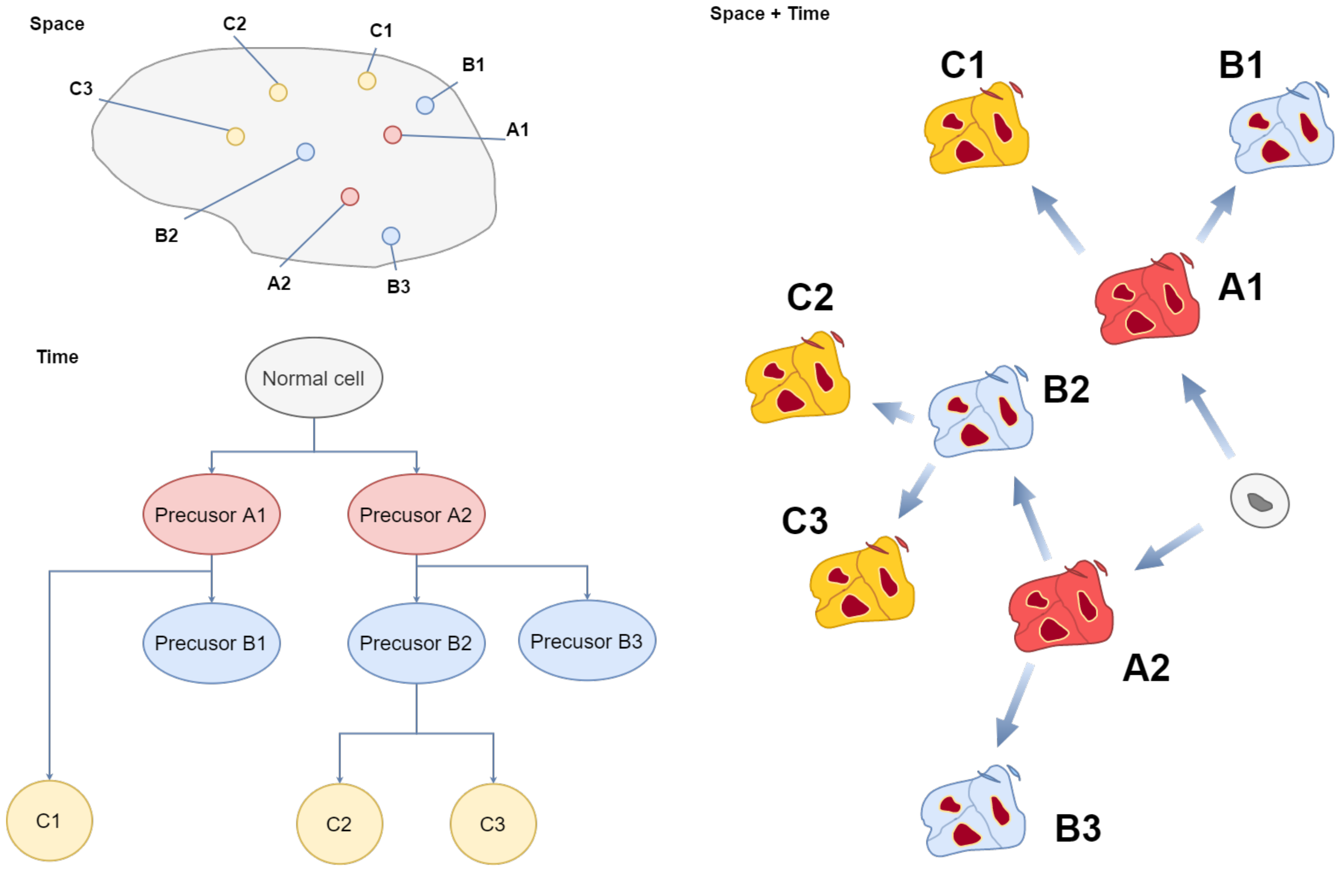


**Supplementary Figure 5.** Tumour progression can be reconstructed in space and time. This example shows the spatiotemporal reconstruction of Glioblastoma progression[2]. The combination of sampling information, reconstructed tumour phylogeny, gene expression profiles, and molecular clock data enables temporal and spatial reconstruction of tumour ontogeny. Together with pseudotime, this concept provided the inspiration for the pseudo-space-time algorithm.

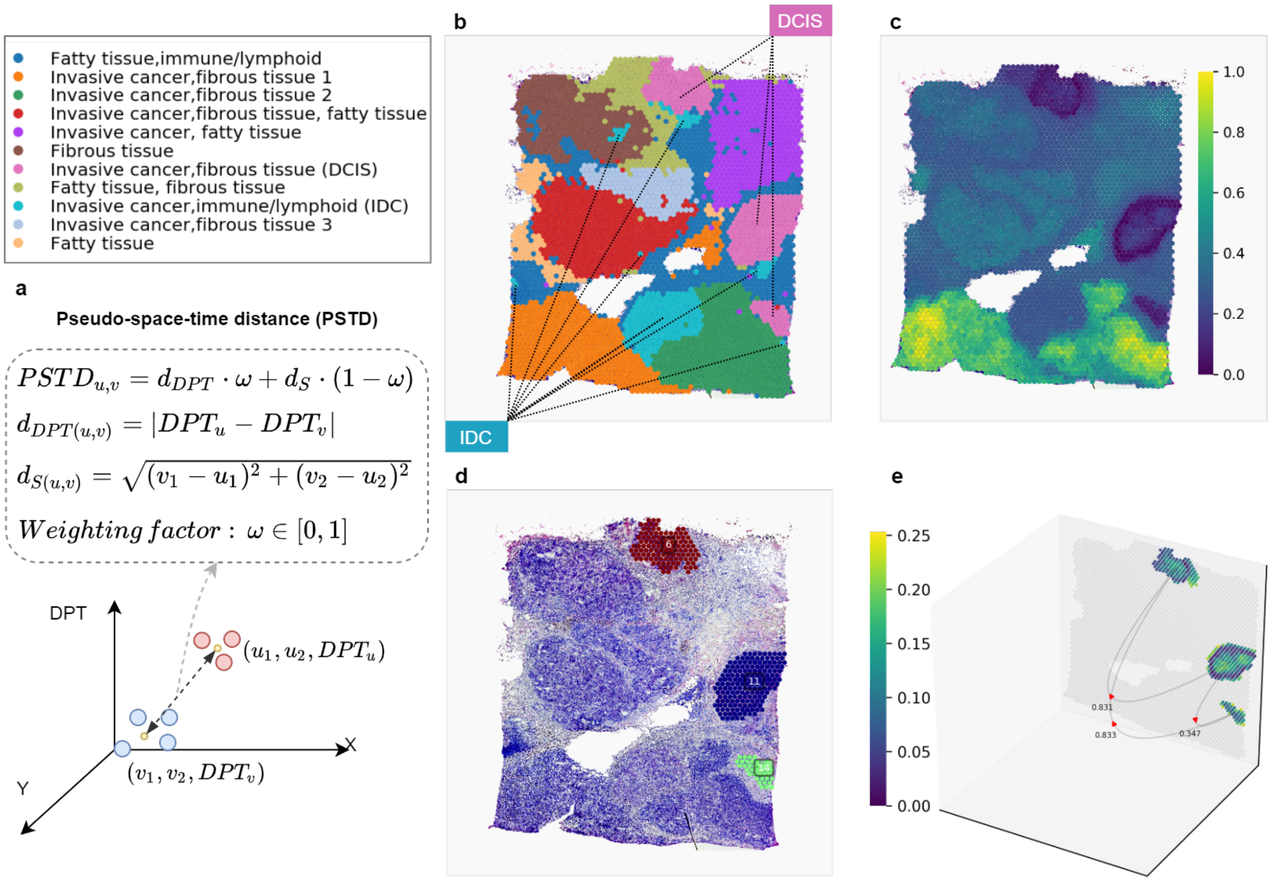


**Supplementary Figure 6.** Spatial trajectory analysis (Case study 3). **a**, A formulae of the pseudo-space-time distance (PSTD). $d_{DPT(u,v)}$ denotes pseudotime distance and $d_{S(u,v)}$ is the spatial distance between two clusters U and V. **b**, Eleven clusters were annotated and labelled based on gene expression patterns derived from PSTD. **c**, Visualization of diffusion pseudotime from gene expression to tissue morphology. The gradient colour scale displays the quantitative general trajectory at the cluster level from the DCIS cluster (lower DPT, 0 as the root) to the IDC cluster (higher DPT, near 1). **d**, The DCIS cluster (pink colour in **b**) is separated into three subclusters by using local clustering based on spatial distance to show the distributed locations of a single cluster within a tissue. **e**, Visualization of the local level of trajectory inference using PSTD.

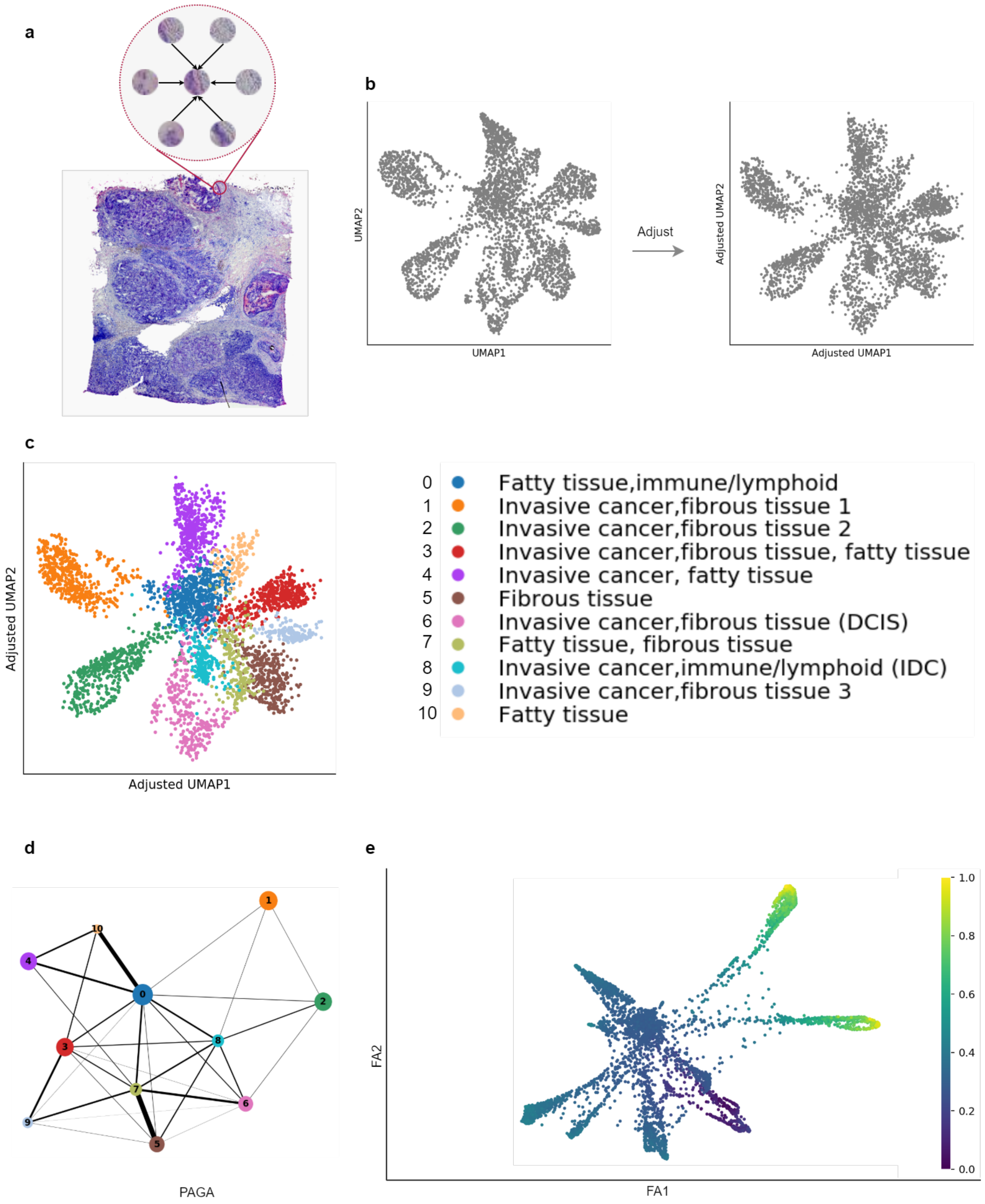


**Supplementary Figure 7.** Analysis of cell types and their relationship in breast cancer dataset. **a**, Adjustment of gene expression levels were carried out by scanning neighbouring spots under the morphological features in the HE image of breast cancer tissue **b**, Visualisation of adjusted gene expression based on UMAP space and spatial distance. **c**, Clustering result in adjusted UMAP space. **d**, PAGA graph showing the relationship among clusters based on gene expression. **e**, Diffusion pseudotime values display in the PAGA embedding space. The gradient colour scale displays the quantitative general trajectory at the cluster level from the DCIS cluster (lower DPT, 0 as the root) to the IDC cluster (higher DPT, near 1), and suggests the likely transition of DCIS-IDC in the tissue.

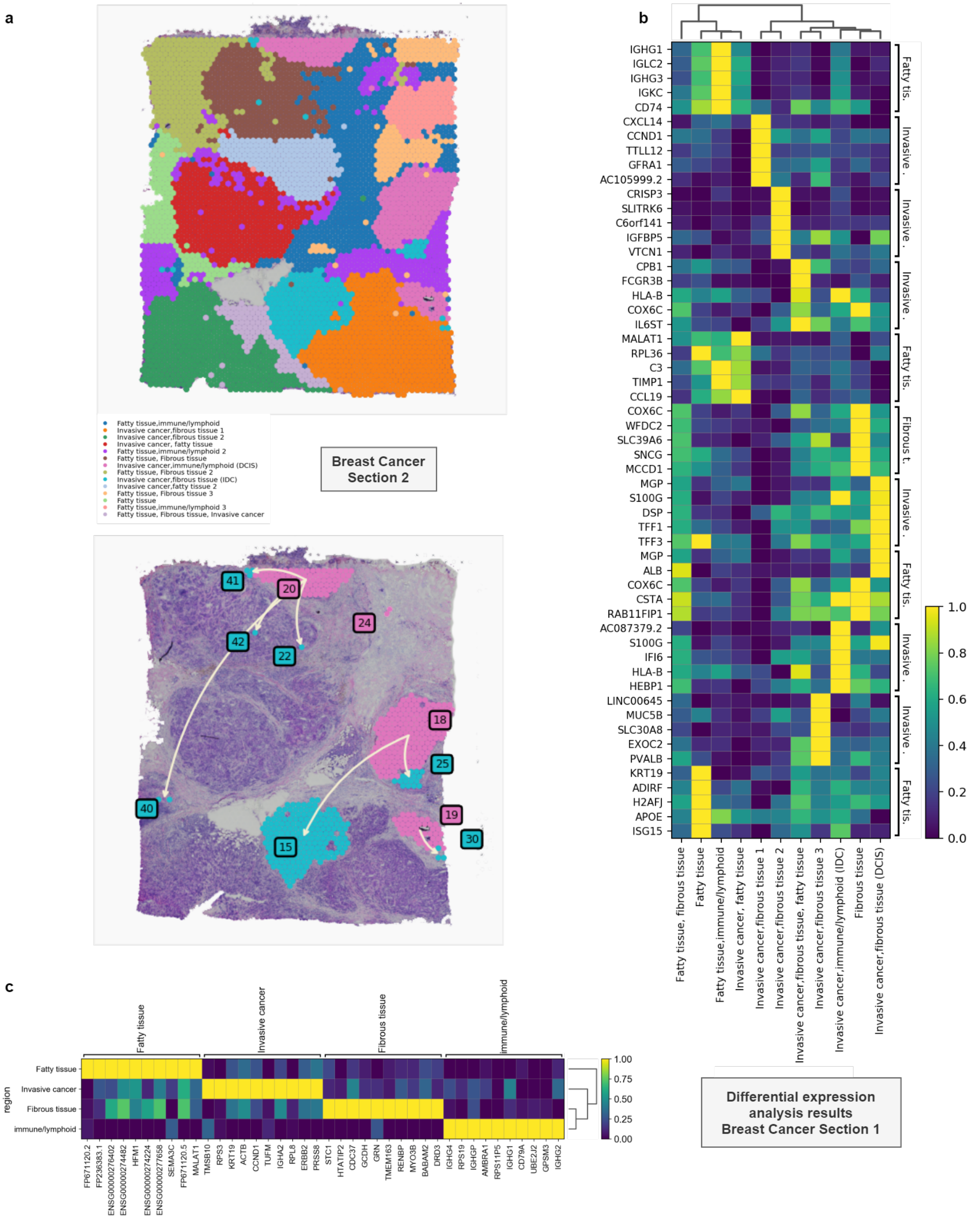


**Supplementary Figure 8.** Cell type annotation of the breast cancer dataset. **a**, SMEclust (top) and PST analysis (bottom) of a ductal carcinoma in situ section-2 dataset. Section-2 was adjacent to the section-1. Analysis results for section-1 are shown in Figure 3 and Supplementary Figures 6, 7, 8b, 8c and 9. **b**, Heatmap of differential gene expression of the whole ductal carcinoma in situ section-1. The marker genes were used for the annotation the clustering results. **c**, Heatmap of differentially expressed genes in a breast cancer dataset generated by High-Density-Spatial-Transcriptomics[3].

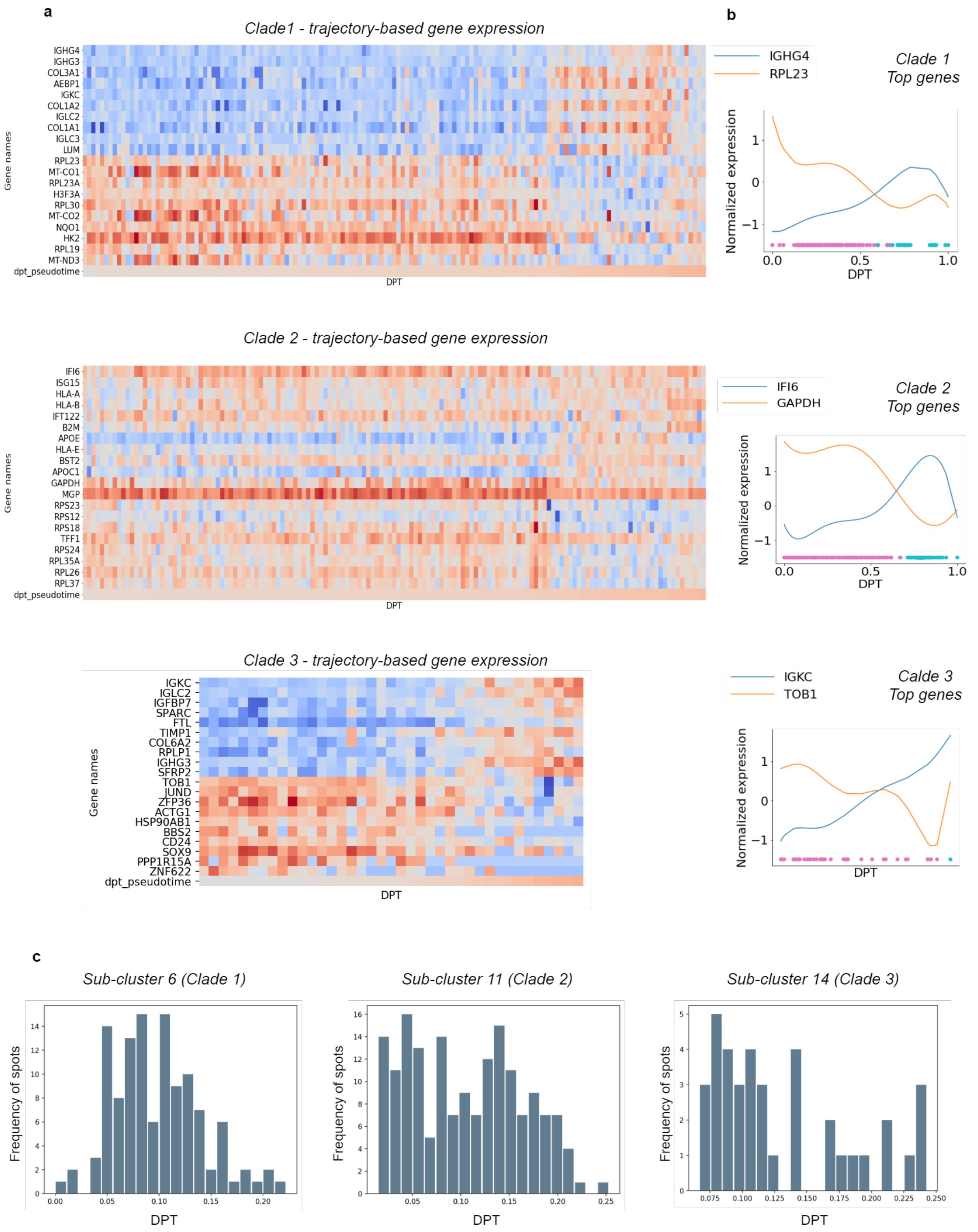


**Supplementary Figure 9.** Analysis of cancer evolution clades identified by PST algorithm. **a**, Heat map of top 10 significant up- and downregulated genes for each clade. **b**, Expression values of single most up- and downregulated gene for each clade during the DPT values transitioning along two clusters. **c**, Histogram of diffusion pseudotime (DPT) values for each clade shows the dynamic distribution of DPT across spots within a clade.

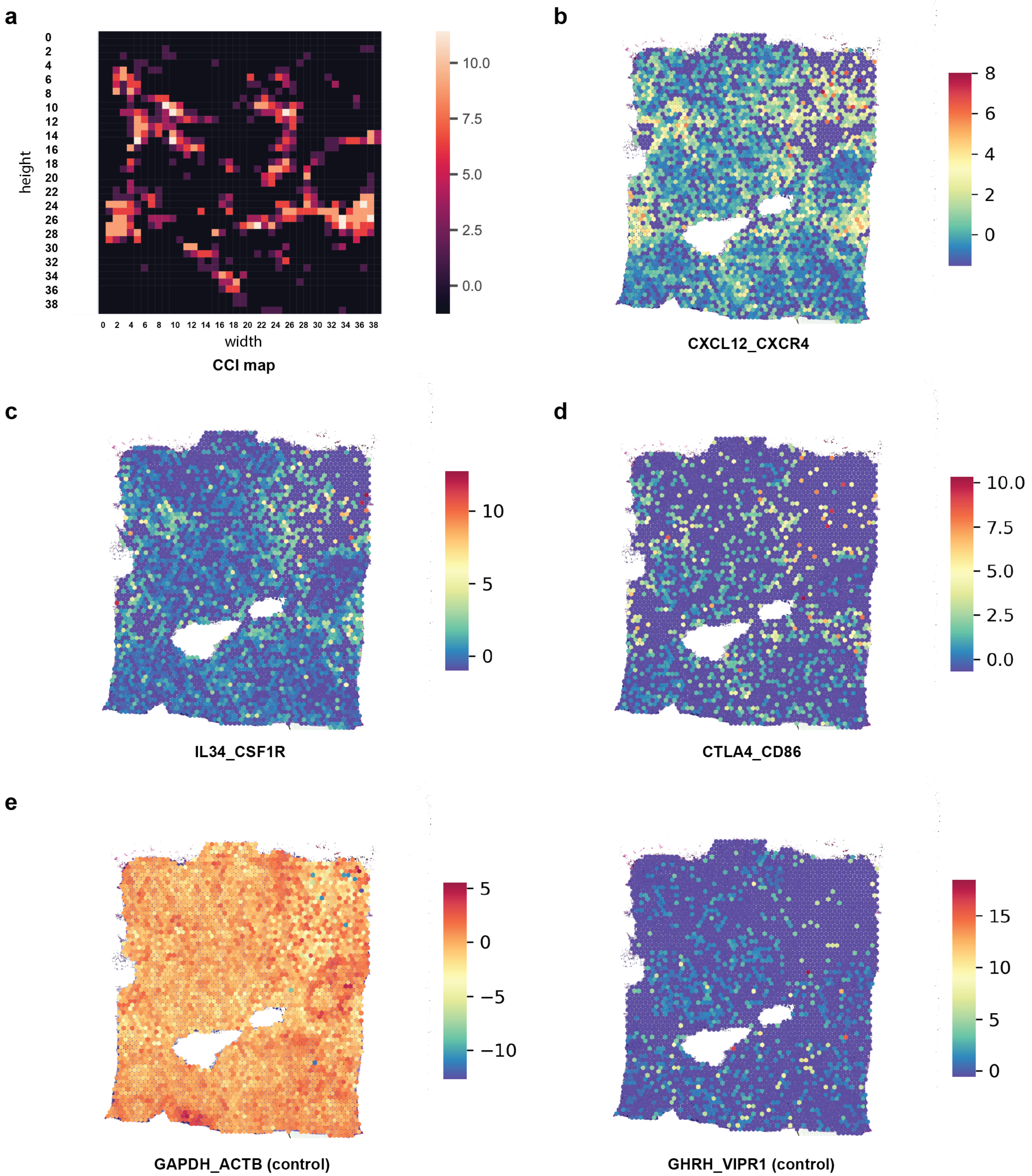


**Supplementary Figure 10.** Comparing CCI map with ligand-receptor expression. The analysis predicts regions with cancer-immune interactions consistent with expression of known ligand-receptor interactions. **a**, Heatmap of CCI showing regions with high ligand-receptor neighbourhood interactions and cell-type diversity. **b-d**, Expression mapping for four ligand-receptor pairs of known immune related interacting genes show similar patterns as CCI enriched regions; **e-f**: two non-interacting control gene pairs show unrelated expression patterns compared to CCI result.

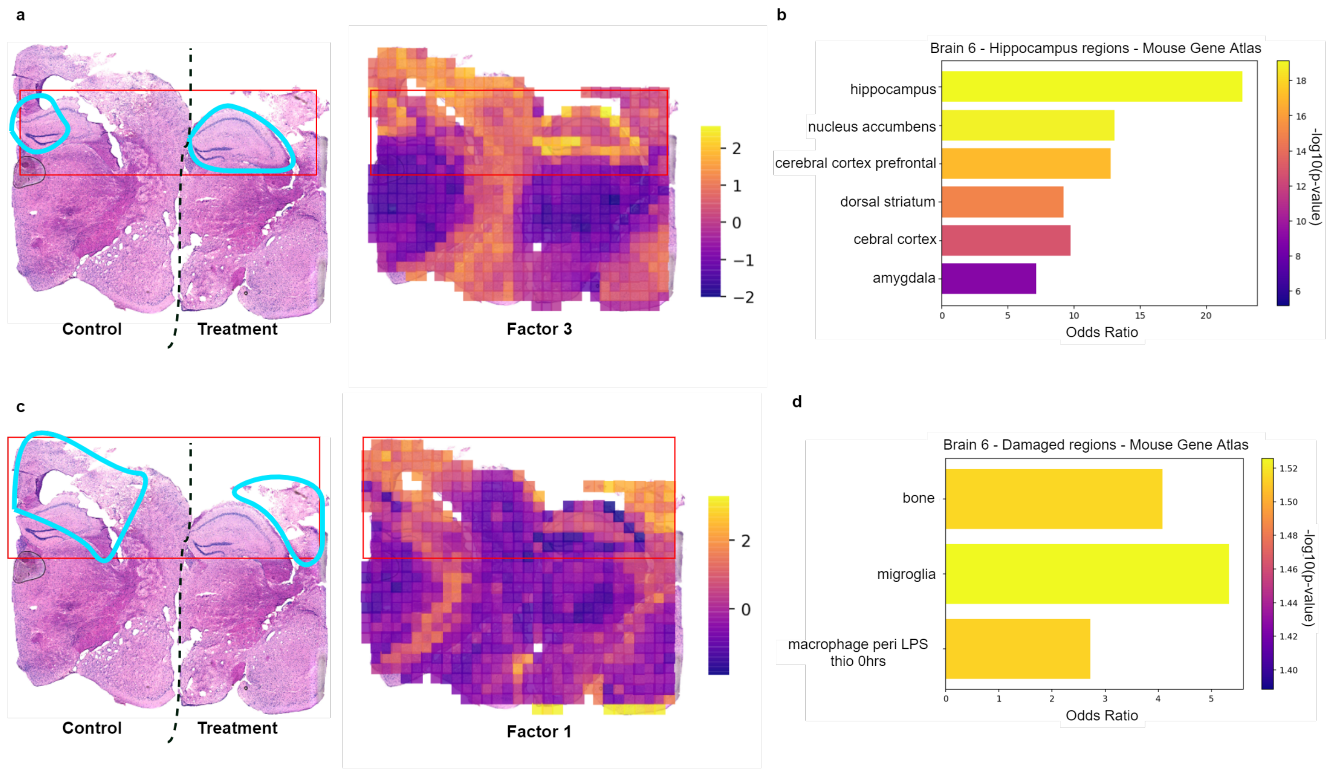


**Supplementary Figure 11.** Factor analysis to find data structure that matches tissue morphological patterns (microenvironments) in damaged brain tissues. An example of a traumatic brain injury mouse model is shown. **a**, Aligning tissue morphology with latent factors. Factor 3 regions have high latent score values and correlate well to hippocampus regions. Two mouse brain sections at the hippocampus regions are shown. The cyan lines indicate hippocampus regions corresponding to the red box in the factor plot. **b**, Enrichment analysis of genes correlated with factor 3 loadings. **c**, Aligning tissue morphology of the tissue damaged regions with factor 1 loadings. Two mouse brain sections are shown. Cyan lines show tissue damaged regions, which align with high loading values for factor 1. **d**, Enrichment analysis result of genes correlated with factor 1.

#### References


**1.** Maynard, K. R. *et al.* Transcriptome-scale spatial gene expression in the human dorsolateral prefrontal cortex. *bioRxiv* DOI: 10.1101/2020.02.28.969931 (2020).

**2.** Sottoriva, A. *et al.* Intratumor heterogeneity in human glioblastoma reflects cancer evolutionary dynamics. *Proc. Natl. Acad. Sci.* **110**, 4009–4014 (2013).

**3.** Vickovic, S. *et al.* High-definition spatial transcriptomics for in situ tissue profiling. *Nat. methods* **16**, 987–990 (2019).
